# A Phosphorus-Limitation Induced, Functionally Conserved DUF506 Protein is a Repressor of Root Hair Elongation in *Arabidopsis thaliana*

**DOI:** 10.1101/2021.07.09.451837

**Authors:** Sheng Ying, Elison B. Blancaflor, Fuqi Liao, Wolf-Rüdiger Scheible

## Abstract

Root hairs (RHs) function in nutrient and water acquisition, root metabolite exudation, soil anchorage and plant-microbe interactions. Longer or more abundant RHs are potential breeding traits for developing crops that are more resource-use efficient and can improve soil health. RH elongation is controlled by both environmental and endogenous factors. While many genes are known to promote RH elongation, relatively little is known about genes and mechanisms that constrain RH growth. Here we demonstrate that a DOMAIN OF UNKNOWN FUNCTION 506 (DUF506) protein, AT3G25240, negatively regulates *Arabidopsis thaliana* RH growth. The *AT3G25240* gene is strongly and specifically induced during P-limitation. Mutants of this gene, which we call *<u>R</u>EPRESSOR OF E<u>X</u>CESSIVE <u>R</u>OOT HAIR ELONGATION 1 (RXR1)*, have much longer RHs, while over-expression of the gene results in much shorter RHs. RXR1 physically interacts with a Rab-GTPase (RXR2), and an *rxr2* mutant phenocopies the *rxr1* mutant. Overexpression of a *Brachypodium distachyon RXR1* homolog resulted in repression of RH elongation in Brachypodium. Taken together, our results reveal a DUF506-GTPase module with a prominent role in repression of RH elongation that is conserved in monocots and dicots.

## INTRODUCTION

Under phosphorus (P) limiting conditions, plants develop shallower primary/basal roots, longer and more lateral roots, longer root hairs (RHs), or cluster roots to improve P foraging and acquisition (Lynch, 2011; Lambers et al., 2015). RHs alone can contribute 70% or more to the total root surface area and can be responsible for up to 90% of phosphate (Pi) uptake (Bates and Lynch, 2001; Jungk, 2001; Haling et al., 2013; Tanaka et al., 2014; Miguel et al., 2015). The combination of long RHs and shallow basal roots in the soil’s P-enriched top layer results in a synergistic effect on P acquisition, and translate into massive increases in biomass in cultivars with both traits (Miguel et al., 2015). Therefore, RHs are potential breeding targets for improving nutrient uptake efficiency in agriculturally important crops (Nestler and Wissuwa, 2016; Rongsawat et al., 2021). RHs are single cell projections from roots that elongate through a process called tip growth. During tip growth, cell expansion is restricted to the cell apex leading to a cell that is cylindrical in shape (Bascom et al., 2018). RHs emerge from specialized root epidermal cells called trichoblasts through a tightly orchestrated series of developmental steps that include RH initiation, tip growth and tip growth termination (Grierson et al., 2014). These steps of RH growth and development are tightly regulated by numerous genes, encoding transcription factors (TFs), and proteins involved in cell wall remodeling, cytoskeletal dynamics and vesicle trafficking. Furthermore, hormones such as auxin, ethylene, jasmonic acid or cytokinin and other signaling molecules including reactive oxygen species (ROS), cytoplasmic Ca^2+^ and phosphoinositides play pivotal roles in modulating RH development (Pitts et al., 1998; Kusano et al., 2014; Mendrinna and Persson, 2015; Mangano et al., 2017; Kato et al., 2019; Han et al., 2020; Tian et al., 2020; Wendrich et al., 2020).

The genetics and molecular mechanisms of RH development have been extensively studied (Parker et al., 2000; Schiefelbein, 2000; Cho and Cosgrove, 2002; Bruex et al., 2012; Lin et al., 2015; Salazar-Henao et al., 2016; Hwang et al., 2017) leading to the identification of a number of key genes. ROOT HAIR DEFECTIVE6 (RHD6), a basic helix-loop-helix (bHLH) TF, plays central roles in RH development (Masucci and Schiefelbein, 1994). The expression of *RHD6* is activated in trichoblasts by R3-type MYB TF CAPRICE (CPC) and its homologs, ENHANCER OF TRY AND CPC1 (ETC1) and TRYPTICHON (TRY), and is suppressed in non-hair cells (atrichoblasts) by plant-specific homeodomain-leucine zipper (HD-Zip) TF GLABRA2 (GL2) (Wada et al., 2002; Menand et al., 2007; Wang et al., 2010; Lin et al., 2015). Both, *CPC* and *GL2*, are transcriptionally regulated by a TF complex composed of the WD40 protein TRANSPARENT TESTA GLABRA1 (TTG1), ENHANCER OF GLABRA3 (EGL3), and the R2R3-type MYB protein WEREWOLF (WER) (Grierson et al., 2014; Slovak et al., 2015). RHD6-LIKE4 (RSL4), which is functionally conserved in higher plants, dominantly regulates expression of various genes involved in RH outgrowth through direct binding to specific *cis*-elements (RH elements; RHEs) in their proximal promoter regions, and consequently controls RH elongation (Yi et al., 2010; Datta et al., 2015; Kim and Dolan, 2016). The MYB TF PHOSPHATE STARVATION RESPONSE1 (PHR1) and its homolog PHR1-LIKE (PHL1), play central roles in the P starvation response of Arabidopsis (Rubio et al., 2001; Bari et al., 2006; Bustos et al., 2010; Rouached et al., 2011; Sun et al., 2016). Overexpression of *PHR1* significantly increases RH length, whereas in the *phr1 phl1* double mutant, RHs are much shorter in P-limiting conditions (Bustos et al., 2010). Conversely, the P-stress responsive TFs WRKY75 and bHLH32 negatively affect RH growth (Chen et al., 2007; Devaiah et al., 2007).Previous studies have also shown that the phytohormones auxin and ethylene synergistically regulate RH growth and differentiation through upregulation of a similar set of genes (Pitts et al., 1998; Bruex et al., 2012; Zhang et al., 2016). Auxin and ethylene reciprocally influence each other’s biosynthesis and distribution, suggesting that complex interactions contribute to developmental outcomes (Růžička et al., 2007; Stepanova et al., 2007). Two models illustrate the roles of auxin and ethylene in the modulation of RH elongation when exposed to low external P (Song et al., 2016; Bhosale et al., 2018). For instance, in P-limiting conditions, the endogenous auxin level is significantly elevated in the root apex, which leads to activation of AUX1-mediated auxin transport and following the induction of ARF19, eventually stimulating RSL2 and RSL4 to enhance RH elongation (Bhosale et al., 2018). On the other hand, ethylene promotes RH growth through transcriptional complexes consisting of EIN3/EIL1 and RHD6/RSL1 as the key regulators of RH initiation and elongation (Feng et al., 2017).

Genetic factors that limit the rate of RH elongation and balance the stimulating effect of auxin, ethylene and RSL2/RSL4 are less well known. The Arabidopsis plasma membrane protein PCaP2with phosphoinositide-binding activity is one such factor (Kato et al., 2019). Also, only few genes involved in termination of RH growth have been identified (Hwang et al., 2016; Shibata and Sugimoto, 2019). Here we show that P-limitation strongly induces a gene encoding a domain of unknown function 506 (DUF506) protein and that it is a strong repressor of RH growth. Overexpression or knockout of the gene results in strong inhibition or stimulation of RH elongation growth, respectively. The gene and its response to P-limitation is conserved in monocot and dicot species. Moreover, the function in RH elongation growth is conserved in Brachypodium, suggesting that knockout or knockdown of DUF506 may be applied to promote RH growth in species of economic value.

## RESULTS

### Expression of AT3G25240 is strongly and specifically induced by P-limitation

*AT3G25240* is a member of a gene family encoding proteins containing domain of unknown function 506 (DUF506). In an RNA-Seq dataset from P-limited Arabidopsis, we found that the *AT3G25240* transcript was massively up-regulated (Fig. S1). The strong induction of *AT3G25240* during P-limitation was confirmed by qRT-PCR, which showed that transcript levels increased nearly 1000-fold following P deprivation and decreased after P re-addition to liquid culture-grown Arabidopsis seedlings (Fig. 1A, B), indicating a direct response of this gene to plant P-status. *AT3G25240* transcript abundance was inversely correlated with media P concentration (Fig. 1C), and induction during P-limitation was attenuated in *phr1* and *phr1*/*phl1* double mutants (Fig. 1D), indicating dependence on these major transcriptional regulators of P-starvation responses. Consistent with this, three PHR1-binding sites (P1BS) are present in the *AT3G25240* promoter (Fig. S2A). In contrast, to the P-limitation response, the response of *AT3G25240* transcript to sulfur, nitrogen or sugar limitation was minor (Fig. 1E). Strong induction of GUS activity during P-limitation in cotyledons and along the main root was detected in *Arabidopsis* seedlings expressing *GUS* under control of the *AT3G25240* promoter (Fig.S2), and GFP fluorescence was evident in P-limited seedlings expressing an AT3G25240-GFP fusion protein under the control of the endogenous promoter (Fig. 1F-M). The 62 kD fusion protein was easily detected by Western blotting in samples from P-limited seedlings (Fig. 1F, left panel). A faint signal was also detectable in samples from P-replete seedlings, but required much longer exposure of the Western blot (Fig. 1F, right panel). During P-limitation GFP signal in live plants was predominantly and highly visible in nuclei of root cells in the RH-forming zone, but not, or much less, in the root tip/expansion zone and the mature, differentiated root (Fig. 1G-J). Closer observation further revealed the presence of the GFP signal in nuclei and the cytoplasm of RH-forming cells (trichoblasts) (Fig. 1K), and in nuclei of differentiated RHs (Fig. 1L, M) and the cytoplasm between nucleus and RH tip (Fig. 1M).

**Figure 1.**
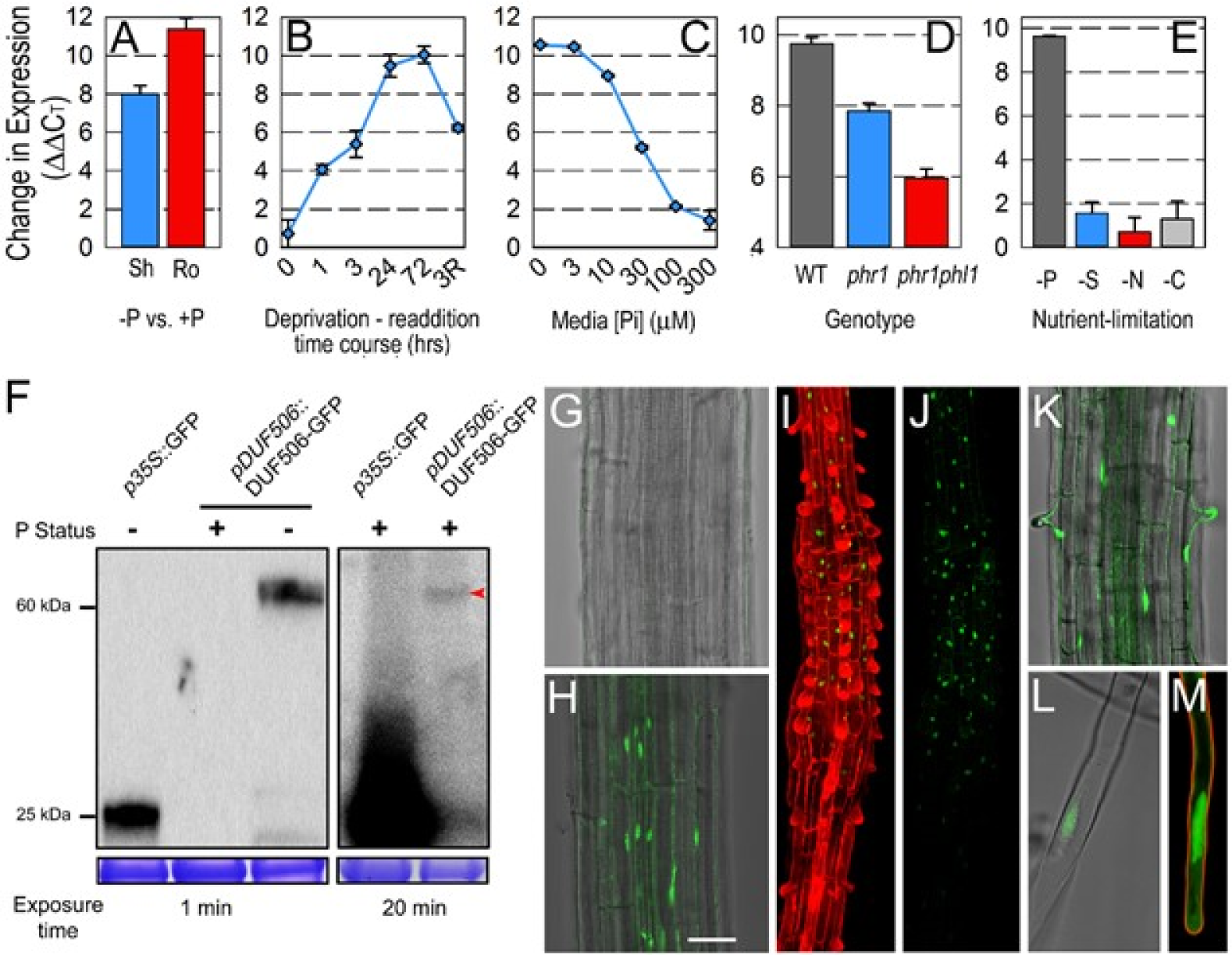
Expression of *At3g25240* transcript and At3g25240protein. (A) Change of *At3g25240*transcript abundance, as measured by qRT-PCR, in P-deprived versus P-replete Arabidopsis seedling shoots (sh) and roots (ro). Fold change is approximately 2^ΔΔCT^ as PCR primer efficiency is close to 100%. (B) Change of *At3g25240* transcript abundance in liquid culture grown Arabidopsis seedlings during a P deprivation / re-addition time course; 3R indicates a 3hrs re-addition of 675µM Pi. (C) *At3g25240* transcript abundance in Arabidopsis seedlings grown on agar plates with various Pi concentrations. (D) *At3g25240* transcript abundance in P-deprived wild type, *phr1* mutant or *phr1phl1* double mutant seedlings, relative to seedlings grown in P-replete conditions. (E) *At3g25240* transcript abundance in P, nitrogen (N), sulfur (S) or carbon (C) -deprived Arabidopsis seedlings. (F) Western blot analysis of P-status dependent abundance of DUF506-GFP fusion protein expressed under control of the endogenous promoter. Coomassie-stained RuBisCO protein is shown as loading control. Short exposure time (1 min, left panel) easily reveals DUF506-GFP in extracts from P-limited (-P) seedlings. Longer exposure time (20 min; right panel) also reveals low DUF506-GFP protein (marked with red arrowhead) in P-sufficient (+P) conditions. (G-M) Detection of GFP signal in roots of Arabidopsis plantlets expressing p*At3g25240::At3g25240*-*GFP*. (G) Absence and (H) nuclear/cytosolic presence of GFP signal (λ=510 nm) in main root under +P and –P conditions, respectively (bar = 50 µm). (I, J) GFP signal in the RH-forming zone of the main root, (K) GFP signal in RH- forming trichoblasts, (L, M) GFP signal in RH nuclei and in the cytoplasm at the RH tip.

### The family of DUF506 proteins

Genes encoding DUF506 proteins are present in monocot and dicot plants species and in green algae, e.g. *Chlamydomonas reinhardtii*, (Fig. S3A), but not in animals. DUF506 proteins harbor two highly conserved 9 and 11 amino acids (aa) motifs (M1, M2) and a conserved ∼80 aa domain (D3) in the C-terminal half, as well as two variable regions at the N- and C-termini (Fig. 2A). In *Arabidopsis thaliana* and *Brachypodium distachyon*, 13 and 9 DUF506 protein-coding genes are annotated, respectively (Fig. 2B and S3A), while in switchgrass (*Panicum virgatum*) and soybean (*Glycine max*) there are 26 and 24, respectively (Fig. S3A). A phylogenetic tree constructed using DUF506 protein sequences from *Arabidopsis thaliana*, *Medicago truncatula, Setaria viridis* and *Brachypodium distachyon* (Fig. S3B) shows clearly defined, higher order branches containing proteins from all four species and sub-branches that reflect the monocot/dicot divide. The response to P-limitation of related DUF506 protein-coding transcripts from different plant species is conserved as well (Fig. S3C).

**Figure 2.**
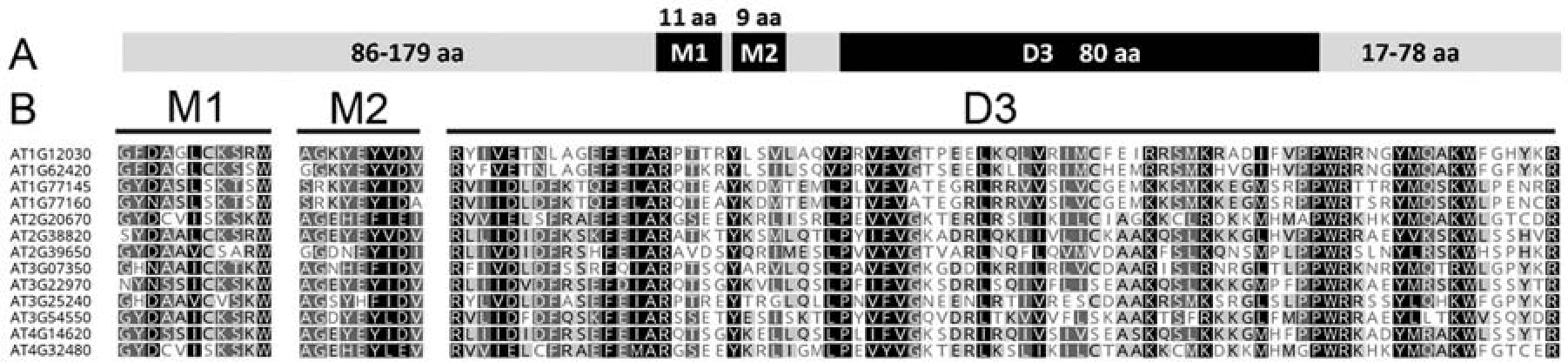
General structure of DUF506 proteins. Graphical depiction of the general one-dimensional structure of DUF506 proteins. Sequences of the conserved motifs (1, 2) and domain (3) in the 13 Arabidopsis DUF506 proteins.

### AT3G25240/RXR1 represses root hair elongation

To explore the biological function(s) of *AT3G25240*, over-expression (OX) lines were generated and a homozygous T-DNA mutant was isolated (Fig.S4). A construct expressing AT3G25240-GFP under control of the *AT3G25240* gene promoter (cf. Fig. 1F-M) was introduced into the T-DNA mutant for complementation, biochemical and subcellular analysis. Roots of 6-day-old OX seedlings displayed shorter RHs while mutant roots had longer RHs when compared to wild type (Fig. 3A). Mutant seedlings expressing AT3G25240-GFP had RHs comparable in length to the wild type, indicating functional complementation, thus linking *AT3G25240* to the long RH phenotype. Quantification of RH length distribution revealed that OX RHs were shorter than 200 µm, with a median length of 65 µm (Fig. 3B). Wild-type RHs ranged from <50 µm to sometimes up to 600 µm, with a median length of 196 µm. RHs of the mutant, subsequently named *rxr1* (*<u>r</u>epressor of e<u>x</u>cessive <u>r</u>oot hair growth* 1), were rarely <100 µm, but ∼17% were longer than 600 µm. The *rxr1* median RH length was 421 µm, i.e more than twice that of wild type. The RH-length distribution plot of the complemented *rxr1*mutant was comparable to wild type. RH growth rates were investigated for *rxr1* mutant, OX and wild type in a single blind study (Fig. 3C). Such analyses revealed that mutant RHs elongate ∼50% faster and OX RHs ∼25% slower than wild-type RHs. The faster RH growth of the mutant was accompanied by a ∼25% higher frequency of cytosolic [Ca^2+^] oscillations (Fig. S5).

**Figure 3.**
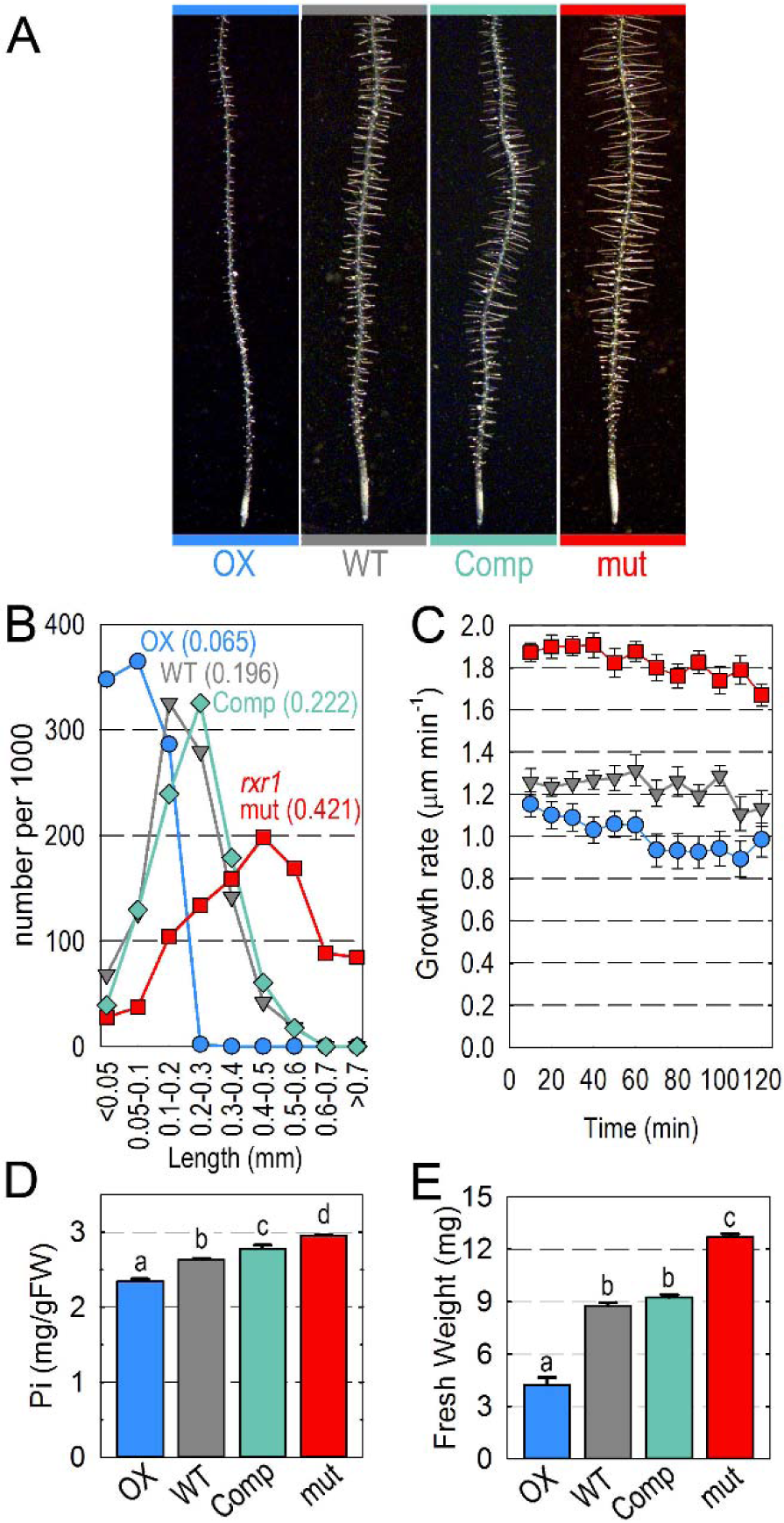
Effect of *RXR1/At3g25240* over-expression or knockout on root hair growth, phosphate content and biomass. (A) RH phenotype and (B) RH length distribution of *RXR1* over-expresser (OX), wild type (WT), *rxr1* mutant (mut) and complemented *rxr1* mutant (Comp). Plantlets were grown on agar (0.8% w/v) plates prepared with nutrient solution containing 675 µM phosphate. For the distribution plots between 787 and 819 root hairs from 10 seedlings were measured for each genotype and numbers for each length category normalized to 1000 root hairs. Numbers in parentheses indicate median values (in mm) for each genotype. (C) Time course of RH growth rate in *RXR1* over-expresser (blue), wild type (grey) and *rxr1* mutant (red). Data represent mean values ± SEM (n = 8).(D) Shoot phosphate concentration and (E) fresh biomass of *RXR1* over-expresser (OX), wild type (WT), *rxr1* mutant (mut) and complemented *rxr1* mutant (Comp) seedlings. Data represent mean values ± SEM (n = 15). Statistical significance of differences was tested by one-way ANOVA analysis (P<0.01) and is indicated by lower case letters.

The various genotypes were also grown in additional conditions and compared with regard to RH length (Fig. S6, S7). Irrespective of P supply or the nature of the medium, the *rxr1* mutant showed the long RH phenotype, while *RXR1* OX RHs were consistently the shortest, and virtually absent in liquid medium. Interestingly, RHs of *rxr1* mutant and *RXR1* OX were still able to respond to P-limitation with an increase in length (cp. Fig. 3A with Fig. S6A, Figs. S6C, D and S7), suggesting that signaling pathways that positively affect RH length during P-limitation are operational. Given the induction of *RXR1* during P-limitation and the importance of RHs for phosphate uptake (Bates and Lynch, 2001), we checked whether shorter and longer median RH length in *RXR1* OX and *rxr1* mutant correlated with seedling phosphate (Pi) content. Indeed, *RXR1* OX seedlings consistently displayed reduced shoot Pi and *rxr1* mutant increased shoot Pi contents (Fig. 3D, Table S2), whereas the complemented mutant had a Pi content more similar to the one of the wild type. RH length also positively correlated with seedling fresh weight (Fig.3E, Table S2) with *rxr1* mutant seedlings being 35% heavier than wild-type seedlings, whereas *RXR1* OX seedlings had a biomass only 50% that of wild type (Fig. 3G). This suggests that longer RHs not only enable higher Pi acquisition but also that higher acquisition results in better growth.

### Repression of root hair-specific gene transcripts in RXR1 over-expressers

Due to the nuclear localization of RXR1, gene-chip transcript profiling was performed with *RXR1* OX, wild type and *rxr1* mutant roots. Although the transcriptional changes in *RXR1* OX and *rxr1* mutant roots compared to wild-type roots were overall minor (not shown), a set of more than 20 gene transcripts previously associated with RHs (cf. Table S3) were found to be slightly (∼2-fold) repressed in *RXR1* OX root samples (Fig. S8; Table S3). No induction of these gene transcripts was found in *rxr1* mutant roots with faster growing RHs, while previous gene-chip profiling of *RSL4* OX roots with long RHs showed 2- to 3-fold induction of many of these transcripts (Yi et al., 2010).

When re-examined by qRT-PCR, the repression of the gene transcripts in *RXR1* OX root samples was confirmed, and ∼2-fold induction of a few of the RH-specific gene transcripts (*PER7*, *RHS15*, *IRT2, At3g07070, At5g04960*) became visible in *rxr1* root samples (Fig. S8). In comparison, *RSL4* OX roots show ∼10-fold induction of *PER7* when examined with qRT-PCR (Yi et al., 2010). Taken together, the results show that RXR1, despite the associated strong RH phenotype, only mildly affects expression of RH-specific gene transcripts, which may suggest that RXR1 also functions in post-transcriptional regulation.

### RXR1 affects root hair elongation independently of auxin or ethylene

Auxin and ethylene are plant hormones known for their role in RH development and elongation (Song et al., 2016; Feng et al., 2017; Bhosale et al., 2018). To test whether RXR1 and auxin (indole acetic acid; IAA) interact during RH elongation, *RXR1* OX, wild type and *rxr1* mutant were grown in ±100 nM IAA (Fig. 4A-D; also cf. Fig.S9A-C for ±10 nM IAA). In all three genotypes, 100 nM IAA significantly stimulated median RH length by 160-170 µm (Fig. 4B-D), indicating that neither loss nor overexpression of *RXR1* affects auxin sensitivity or perception. 100 nM IAA or knockout mutations in major genes (*RSL4*, *ARF19*, *AUX1*, *TAA1*) involved in RH auxin signaling (Bhosale et al., 2018) also did not affect *RXR1* transcript abundance in seedlings grown ± phosphate (Fig. S9G, H). Furthermore, the expression of well-characterized auxin signaling pathway genes in *RXR1* OX or *rxr1* mutant was not affected (Table S4), leading to the conclusion that RXR1 acts independently of auxin in modulating RH elongation. Similar results were obtained with ethylene (Fig. 4E-H, Fig. S9D-F, I-K). Addition of 1 µM 1- aminocyclopropane-1-carboxylic acid (ACC) to Gelzan plates increased median RH length strongly in OX, wild-type and *rxr1* mutant (261, 229 and 198 µm respectively; Fig. 4F-H), and 1 µM ACC or mutations in *EIN3* or *CTR1* did not affect *RXR1* expression (Fig.S9I, J). Moreover, OX and *rxr1* mutant also did not differ from wild type in the classical response of dark-grown hypocotyls to increasing ACC doses (Fig. S9K). Furthermore, well-characterized ethylene signaling pathway genes were unaffected in *RXR1* OX or *rxr1* mutant (Table S4).

**Figure 4.**
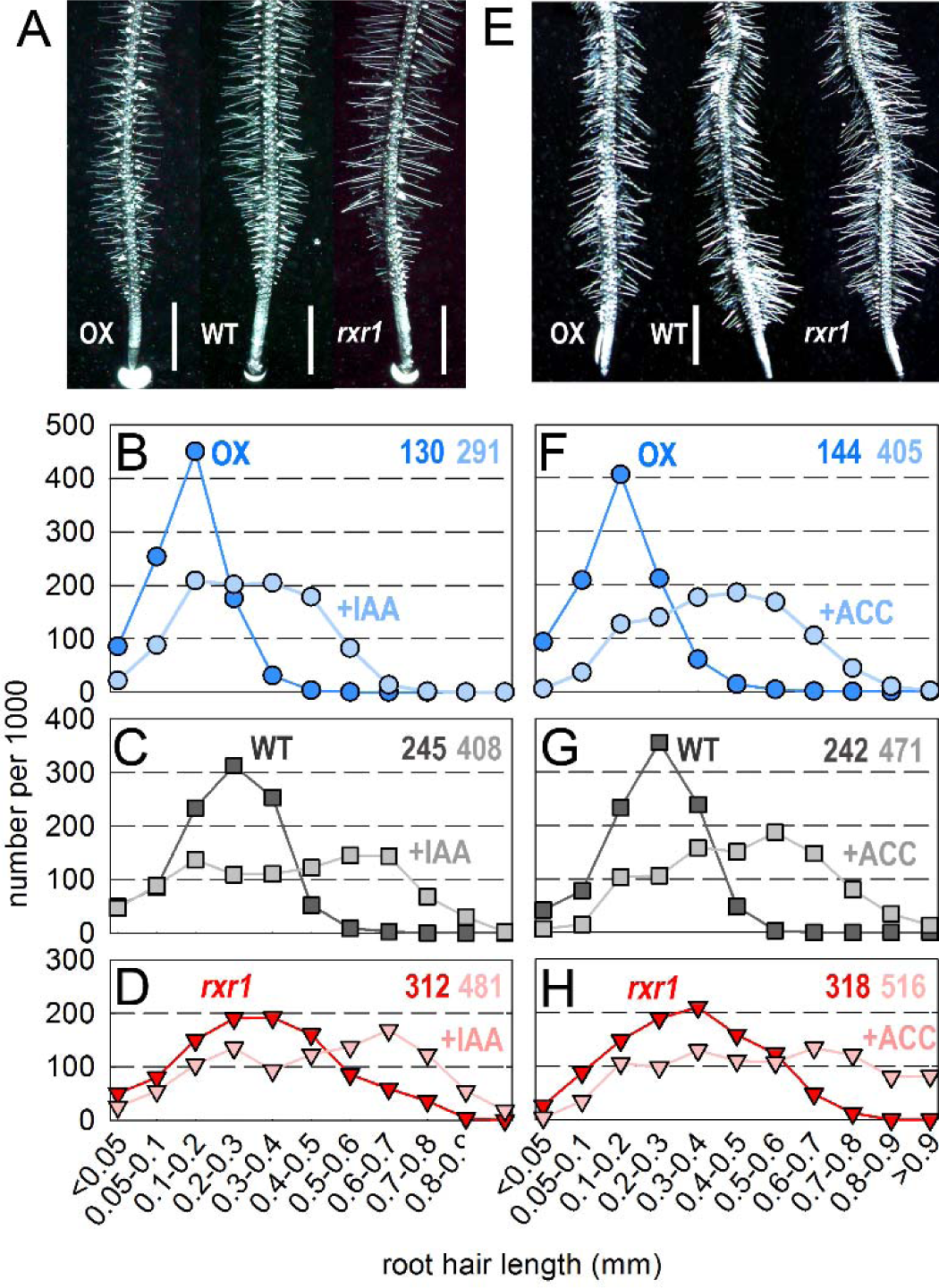
Effect of auxin and ethylene on RH length of *RXR1* over-expressers and *rxr1* mutants. Effect of (A-D) 100 nM IAA or (E-H) 1 µM ACC on root hair length of *RXR1* over- expresser, wild type and *rxr1* mutant. (A, E) RH phenotype of seedlings grown on Gelzan (0.4% w/v) plates prepared with nutrient solution containing 675 µM phosphate and (A) 100 nM IAA or (E) 1 µM ACC (bars = 1 mm). (B-D, F-H) RH length distribution plots for *RXR1* over-expresser (OX), wild type (WT), and *rxr1* mutant (*rxr1*) grown in the absence (darker colors) or presence of (B-D) 100 nM IAA or (F-H) 1 µM ACC (lighter colors). For the distribution plots between 571 and 958 root hairs from 10 seedlings were measured per genotype and treatment, and numbers for each length category were normalized to 1000 root hairs. Numbers in the top right corners indicate color-coded median values (in µm).

### RXR1 interacts with a Rab GTPase

To elucidate a possible post-transcriptional mode-of-action for RXR1 and to identify RXR1-interacting proteins, two independent immuno-precipitation (IP) experiments followed by mass-spectrometric (MS) analysis of tryptic protein digests were conducted using the RXR1-GFP complemented *rxr1* line (cf. Figs.3, S4). Table 1 summarizes the proteins identified in both experiments (cf. Table S5 for more details). A Rab GTPase (RabD2c/AT4G17530) showed the highest sequence coverage. Because of the known roles of Rab GTPases in directional growth processes (Preuss et al., 2004; Szumlanski and Nielsen, 2009; Peng et al., 2011), the interaction of RXR1 and RabD2c was investigated further *in vitro* and *in vivo* (Fig. 5). GFP-tagged RXR1 protein, expressed under its native promoter, was successfully immuno-precipitated in samples generated from P–limited seedlings using recombinant His6-tagged RabD2C protein and Anti-His6 antibody (Fig. 5A). Similar to previous results (cf. Fig. 1F) no RXR1-GFP protein was detectable after short (1 min) exposure in samples prepared from P-sufficient seedlings, nor was GFP itself immuno-precipitated by His6-tagged RabD2C and Anti-His6 antibody (Fig. 5A).

**Figure 5.**
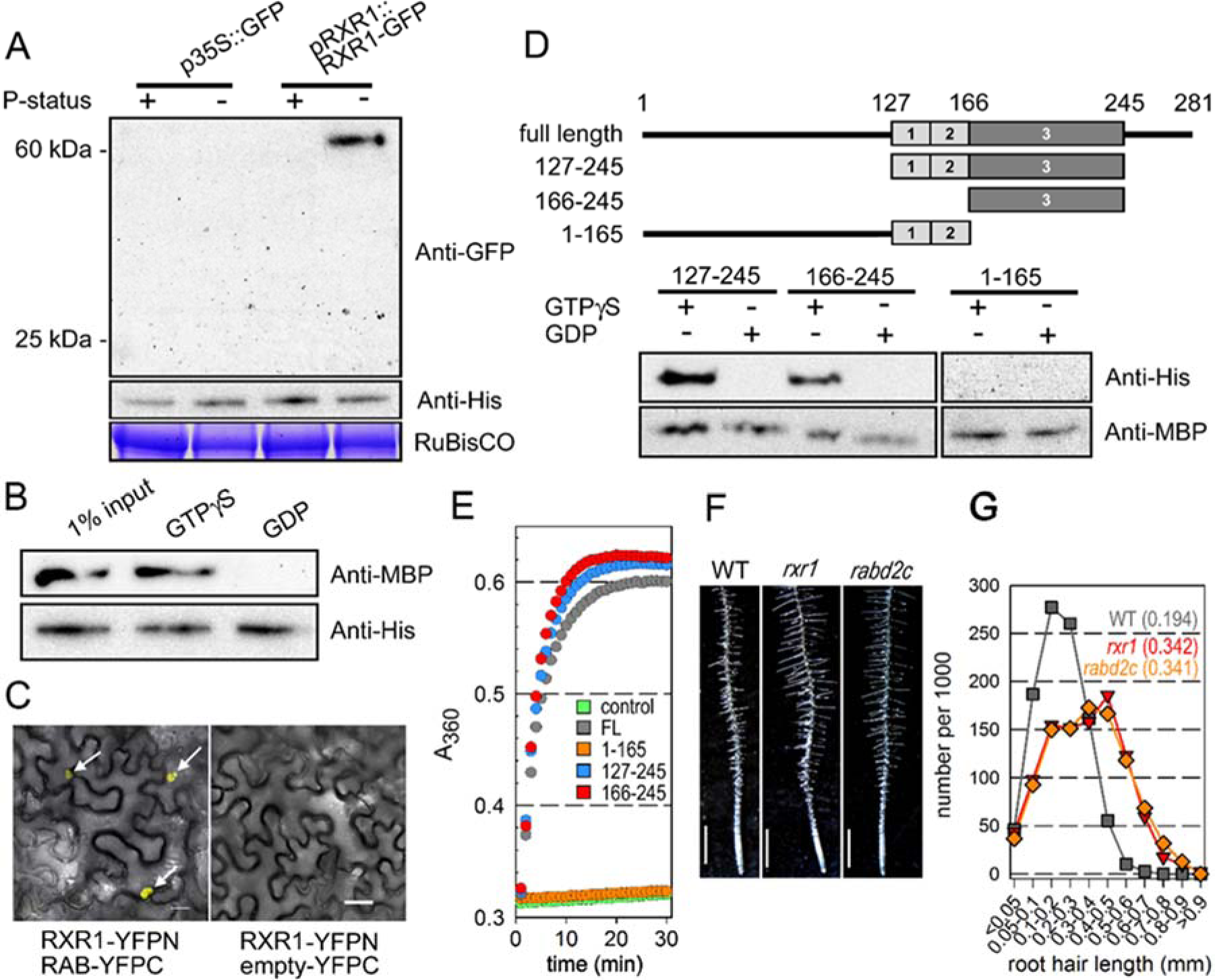
Interaction of RXR1/At3g25240 and Rab GTPase/At4g17530. (A) Immuno-pull down of 62 kD RXR1-GFP fusion protein expressed under control of the endogenous promoter (cf. Fig. 1F) with His6-tagged Rab GTPase and Anti-His antibody (right lane). Exposure time of the Western blot after treatment with anti-GFP antibody conjugated to horseradish peroxidase was 1 min. (B) Immuno-pull down of MBP-tagged DUF506 protein with His-tagged Rab GTPase and Anti-His antibody in the presence of GTP but not GDP. (C) Bimolecular fluorescence complementation in *Nicotiana benthamiana*. Arrows point to nuclei showing yellow fluorescence as a result of RXR1-YFPN and RAB-YFPC interaction. (D) Immuno-pull down of His-tagged Rab GTPase with partial/truncated MBP-tagged RXR1 proteins and Anti-MBP antibody. (E) Rab GTPase activity assay in the absence or presence of the four RXR1 protein versions depicted in (D). (F) RH phenotype (bars = 1 mm) and (G) RH length distribution plots of wild type, *rxr1* and *rabd2c* mutants grown on agar plates as in Figure 3A.

**Table 1.** RXR1-interacting proteins.

| AGI | Annotation | Molecular mass (kD) | Unique peptides (IP1 / IP2) | Coverage (%) (IP1 / IP2) |
| --- | --- | --- | --- | --- |
| At1g12000 | Phosphofructokinase | 61 | 3 / 3 | 8.0 / 8.1 |
| At1g64440 | UDP-Glucose 4-epimerase | 38 | 3 / 2 | 8.9 / 7.2 |
| At1g74960 | Beta-Ketoacyl-ACP Synthetase 2 | 58 | 2 / 3 | 3.9 / 7.0 |
| At1g78850 | Mannose binding lectin 1 | 49 | 1 / 2 | 3.0 / 6.6 |
| At2g46520 | Exportin-2 | 109 | 6 / 4 | 7.3 / 5.5 |
| At3g05970 | Long-chain acyl-CoA synthetase 6 | 77 | 5 / 6 | 10.1 / 12.7 |
| At3g62530 | ARM repeat superfamily protein | 25 | 4 / 3 | 18.6 / 16.7 |
| At4g17530 | GTPase homolog RabD1C | 22 | 2 / 4 | 11.4 / 30.2 |
| At4g23850 | Long-chain acyl-CoA synthetase 4 | 75 | 3 / 5 | 5.7 / 7.7 |
| At5g11720 | Alpha-glucosidase | 101 | 3 / 4 | 4.0 / 5.6 |
| At5g42150 | Glutathione S-transferase | 36 | 2 / 7 | 5.1 / 25.7 |

RXR1-GFP was immuno-precipitated with GFP antibodies from roots of P-limited plants expressing *pRXR1:RXR1*-GFP, and co-precipitated proteins were identified by mass spectrometry. Two independent experiments (IP1, IP2) were performed. Proteins identified in samples from RXR1-GFP expressing plants in both experiments, but not in samples from plants expressing GFP, are listed. Using BLAST-P, protein identification probability for each protein was 100% in each of the two experiments. AGI: Arabidopsis Gene Identifier. Coverage (in %) is defined as the number of amino acids covered by the identified peptides divided by the total number of amino acids of the protein.

Rab GTPases are active and interact with specific effector proteins in the GTP-bound state (Stenmark, 2009). Hydrolysis of GTP converts the active back to the inactive conformation and leads to dissociation of the effector protein(s). To test GTP-dependency of RXR1-RabD2c interaction, both proteins were produced in bacteria with maltose-binding protein (MBP) and His6-tags, respectively, and incubated together either in the presence of GDP or GTPγS, followed by IP with Anti-His6 antibody and Western detection with Anti-MBP antibody. RXR1-RabD2c interaction was undetectable in the presence of GDP, but was evident with GTPγS (Fig. 5B). RXR1 and RabD2c also interacted *in vivo* in bimolecular fluorescence complementation (BiFC) assays following infiltration of RXR1-YFPN and RabD2c-YFPC into *Nicotiana benthamiana* leaves. Yellow fluorescence, indicative for interaction, was present in nuclei (Fig. 5C; left). No yellow fluorescence was present when RXR1-YFPN and YFPC were co-injected (Fig. 5C; right). In addition, full-length and three truncated variants of RXR1 were tested for *in vitro* interaction with and activity of RabD2c (Fig. 5D, E). These experiments showed that interaction and GTPase activity are both dependent on RXR1 domain 3 (cf. Fig. 2A).

The RXR1-RabD2c interaction was also consistent with the agreement of *rxr1* and *rabd2c* mutant RH length and length distribution phenotypes (Fig. 5F, G; Fig.S10), hence *RabD2c* was named *RXR2*. Moreover, RabD2c/RXR2-GFP fusion protein complemented the RH phenotype of *rabd2c/rxr2*, and displayed an expression pattern similar to RXR1- GFP (Fig. S11; cp. to Fig. 1M, L). Taken together the results show that a RXR1- RabD2c/RXR2 module negatively regulates RH elongation.

### Conserved regions of RXR1 are not sufficient to inhibit root hair elongation growth

Since amino acids 127-245 of RXR1, comprising the conserved motifs 1/2 and domain 3, were sufficient for interaction with and activity of RabD2c/RXR2 (Fig. 5D, E), their biological function was further investigated by transforming Arabidopsis wild type with RXR1127-245 -GFP driven by the CaMV35S promoter. Over-expression of the 46 kD fusion protein was detected in stably transformed lines (Fig. S12A) and localization of RXR1127-245 -GFP in RH nuclei (Fig. S12B) was similar to full length RXR1-GFP. However, over-expression of truncated RXR1127-245 no longer reduced RH elongation growth (Fig. S12C, D). Similarly, seedling biomass and Pi content of RXR1127-245 OX lines was not reduced (Fig. S12E, F). The construct was also introduced into *rxr1*, but did not complement the long RH phenotype (Fig. S12C). These results indicate that elements outside of the conserved motifs/domain of RXR1 are required to reduce RH elongation.

### Investigation of additional RXR1 interactors

To find out whether RXR1 interacts with other identified candidates (Table. 1), we continued to isolate and phenotypically characterize four additional T-DNA mutants, which have been demonstrated the disruption of their donor genes. No distinguishable RH length changes were observed from SALK_111594/*At1g12000* and CS459880/*At1g78850* mutants, whereas similar longer RH phenotypes were respectively noticed from two LACS (<u>L</u>ong-chain <u>a</u>cyl-<u>C</u>oA <u>S</u>ynthetase) mutants, LACS4/At4g23850 and LACS6/At3g05970 (Fig. 6A, B). Both of them exhibited at least 50% (+P) or 28% (- P) longer RH length in relative to wild type (Fig. 6C, D), suggesting that RXR1 might synergically associate with LACSs while interacting with RabD2c (Fig. 8).

**Figure 6.**
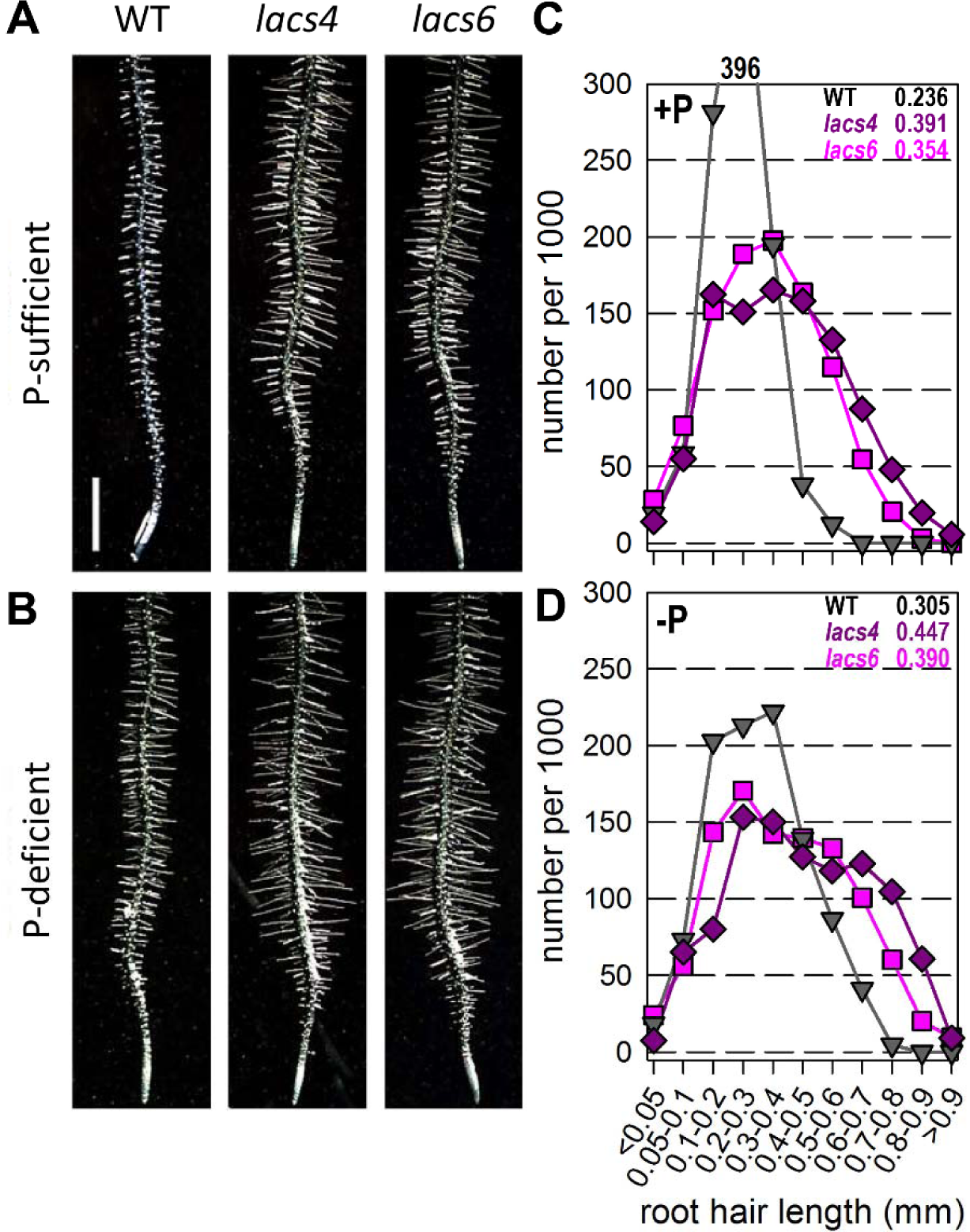
RH length phenotypes of *lacs4* and *lacs6* mutants. LACS4 (At4g23850) and LACS6 (At3g05970) were identified as RXR1 interactors (cf. Tables 1 and S5). Corresponding homozygous T-DNA mutants (*lacs4*, CS65779; *lacs6*, CS2103801) and wild type were grown on (A) P-sufficient or (B) P–deficient agar plates to assess RH length differences. (C, D) RH length distribution of wild type (grey), *lacs4* (purple) and *lacs6* (pink) mutants grown on Gelzan (0.4% w/v) plates prepared with nutrient solution containing (C) 675 µM or (D) no phosphate for 3 days. Between 659 and 878 RHs were measured for each genotype and numbers for each length category normalized to 1000 RHs. Numbers in the top right corner indicate color-coded median values (in mm) for each genotype.

### A close RXR1 homolog reduces root hair elongation in Brachypodium

*AtRXR1* has close P-limitation induced homologs in other species (cf. Fig. S3). To test if the closest *Brachypodium* homolog (*Bd2g58590*), besides being P-limitation induced (cf. Fig. S3C), also affects RH length, a construct over-expressing *Bd2g58590* fused to *GFP* was infiltrated into *Nicotiana benthamiana* leaves. Similar to AtRXR1, BdRXR1-GFP localized to the nucleus (Fig. 7A). Stable OX lines were then generated in *Brachypodium* leading to 50-100 fold higher expression levels as indicated by qRT-PCR analysis (Fig. 7B; ΔΔCT = 7-8). The overexpressing *BdRXR1* lines were analyzed for their RH phenotype. Similar to overexpression of *AtRXR1*, overexpression of *Bd2g58590* resulted in reduced RH length (Fig. 6C, Fig. S13), while RH density and main root diameter (Fig. 7D) were not noticeably changed. The RH length and 2-dimensional RH convex envelope of the OX1 line was 50% reduced compared to wild type (Fig. 7E, F) suggesting that OX1 roots have a much smaller (<25%) rhizosphere than the wild type. Also similar to the results with *AtRXR1* OX, over-expression of *Bd2g58590/BdRXR1* in *Brachypodium distachyon* resulted in reduced plant biomass production (Fig.7G, H). Based on these results we conclude that *Bd2g58590* and *AtRXR1* are orthologous genes.

**Figure 7.**
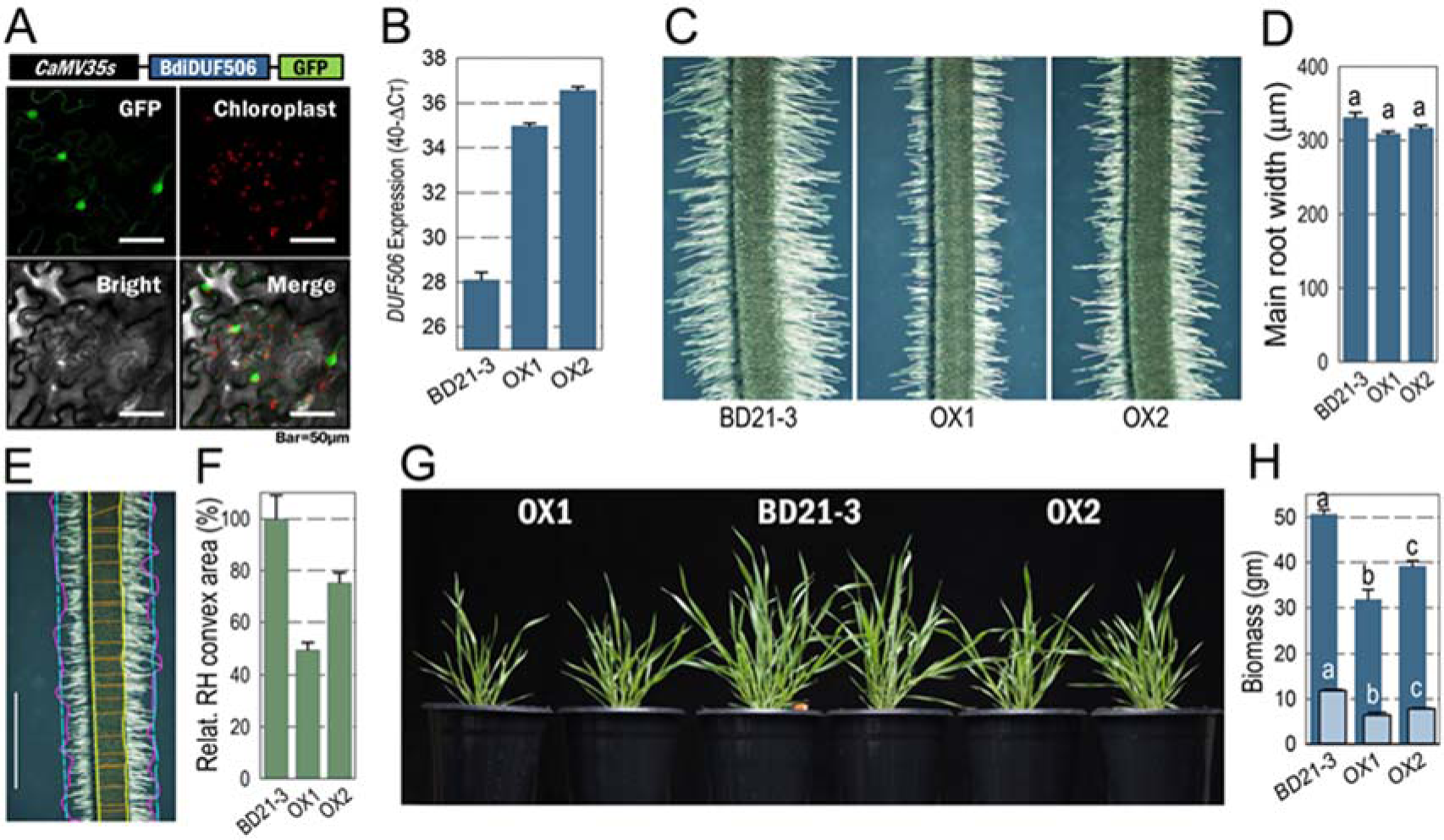
Overexpression of a close *RXR1* homolog in *Brachypodium distachyon*. Sringe-infiltration of a p*CaMV35S*::*BdDUF506*-*GFP* construct into *Nicotiana benthamiana* leaves results in nuclear localization of the fusion protein (bar = 50 µm). Abundance of *BdDUF506* transcript in stable over-expression lines of *Brachypodium distachyon*. (C) RH phenotypes of *Brachypodium* wild type (BD21-3) and two stable over-expression lines. (D) Quantification of main root width, (E) example for determination of RH convex envelope as defined by the sum of areas between the yellow and cyan lines. The cyan lines are the averages of the magenta lines that connect RH tips (bar = 1 mm).(F)Relative RH convex envelope area (WT = 100%) for the genotypes shown in (C). (G) Growth aspect of the genotypes shown in (C) at an age of six weeks, and (H) shoot fresh and dry weights of the genotypes at an age of eight weeks. Data in (D) and (H) represent mean values ± SEM (n = 10).Statistical significance of differences was tested by one-way ANOVA analysis (p<0.01) and is indicated by lower case letters.

## DISCUSSION

In this work, we identified the DUF506-protein RXR1 and its interacting RabD2c GTPase/RXR2 as repressors of RH elongation in Arabidopsis (cf. Fig. 8) as well as *Brachypodium distachyon*. By doing so, we attributed biological functions to the first DUF506-protein coding genes as well as a not well characterized Rab GTPase. *RXR1* was chosen for research because of its transcriptional response to P-limitation, the presence of potential orthologs in dicot and monocot plant species (Fig. S3), and the general lack of information for the *DUF506* gene family.

**Figure 8.**
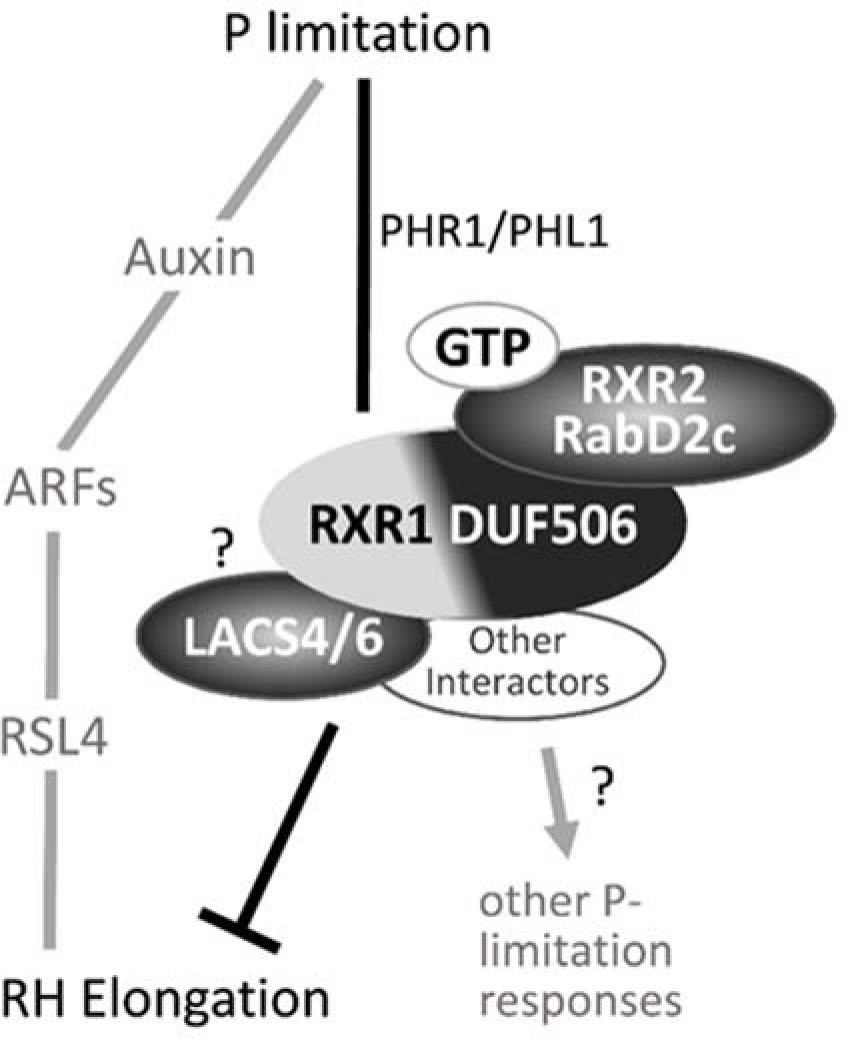
Working model for molecular function of RXR1/RXR2. RXR1/RXR2 is a novel module that represses RH elongation independently of the stimulating effect of the phytohormone auxin and its downstream signaling pathway (cf. (Bhosale et al., 2018). When P is limiting, PHR1/PHL1-dependent *RXR1* expression and protein strongly increase. RXR1 interacts with GTP-bound RXR2 through its conserved DUF506 domain (indicated in black), whereas RXR1-specific sequences (in gray) are required to inhibit RH elongation growth. Biochemical and genetic evidence (i.e. RH growth phenotypes) further suggest RXR1 interaction with LACS4/6, possibly through RXR1-specific sequences. RXR1 interaction with other proteins may modulate additional P-limitation responses.

The discovered interaction of RXR1 and RabD2c/RXR2 spurred subsequent work because RabD2c and other related small GTPases are known to affect pollen tube tip growth (Szumlanski and Nielsen, 2009; Peng et al., 2011; Wang et al., 2012), or localize to tips of growing RHs (Preuss et al., 2004). Pollen tubes and RHs are similar in terms of their highly polarized “tip growth” where cell expansion occurs at the very apex of the cell, and is supported by a tip-focused delivery of membrane and cell wall components (Szumlanski and Nielsen, 2009). RabD2c/RXR2 is highly similar to RabD2b (At5g47200), and both Rab GTPases were found to have overlapping functions in vesicle trafficking during pollen tube growth (Peng et al., 2011). A role for RabD2c/RXR2 in RH development was unknown until now, however.

For the interaction with and activity of RabD2c/RXR2 in the presence of GTP, the conserved DUF506 regions of RXR1 are required and sufficient (Fig. 5, cf. Fig. 8), suggesting that other DUF506 proteins may interact, as so-called effector proteins (Nielsen, 2020), with Rab GTPases in a GTP-dependent manner as well. The variable RXR1 sequences are required for the ability of RXR1 to repress RH elongation, supposedly through interaction with other proteins (see below), eventually resulting in altered cytosolic [Ca^2+^] oscillations (Fig. S5) and RH elongation growth. Our experiments also showed that the RXR1-RXR2 module acts independently of auxin (and ethylene) in controlling RH elongation, with the former exerting negative control and the latter exerting positive control. Therefore, crosstalk or convergence of the auxin and RXR1-RXR2 pathways during RH elongation appear to occur only further downstream (Fig. 8).

The extraordinary response of *RXR1* transcript to P-limitation, its dependence on PHR1/PHL1 (Fig. 1; Fig. 8) (Bustos et al., 2010), and lack of response to other environmental and developmental factors observed in RNA Seq databases such as eFP browser (Winter et al., 2007) or GENEVESTIGATOR (https://genevestigator.com/), suggests an important and specific role for RXR1 during P-limitation. However, despite almost undetectable transcript, RXR1 protein is present at a low level and functional in P-replete conditions, as indicated by the long RHs of P-fed *rxr1* mutants (Fig. 1; Fig. 3). More RXR1 protein may be needed during P-limitation to counteract a strong inducing effect of auxin (Bhosale et al., 2018) and balance RH elongation growth. RXR1 may also possess other biological functions, as suggested by its spatial expression during P-limitation (Fig. S2) and by the existence and nature of RXR1-interacting proteins (Table 1). In this context, T-DNA mutants of *At1g12000* and *At1g78850* (cf. Table 1) did not show long RH phenotypes, suggesting biological roles unrelated to RH growth, whereas mutants of *At3g05970* and *At4g23850* (cf. Table 1), encoding long-chain acyl-CoA synthetases (LACS4, LACS6), displayed *rxr1*-/*rxr2*-like RH phenotypes (Fig. 6). RXR1/RXR2, through interaction with peroxisomal LACS6 (Fig. 8), therefore may limit beta-oxidation (Fulda et al., 2002; Shockey et al., 2002)and thus energy production required for RH elongation growth. On the other hand, interaction with LACS4 could indicate that some other aspect of RH lipid metabolism, such as the provision of activated fatty acids for membrane lipid or phosphoinositide synthesis, is controlled by RXR1/RXR2.During RH tip growth, membrane lipids are required in large amounts for the expanding plasma membrane, while phosphatidylinositol 4,5-bisphosphate (PtdIns(4,5)P2) is a key signaling molecule in the process(Stenzel et al., 2008; Kato et al., 2013; Kusano et al., 2014; Kato et al., 2019) Despite the absence of mutant RH phenotypes, the aforementioned RXR1-interacting proteins At1g12000 (β-subunit of pyrophosphate-dependent phosphofructokinase; PPi-PFK) and At1g78850 (mannose-binding lectin1) are intriguing. PPi-PFK is a key enzyme that facilitates glycolytic flux during severe Pi stress when the intracellular levels of adenylates and Pi are very low (Plaxton and Tran, 2011), whereas At1g78850 enhances purple acid phosphatase 26 (PAP26) function to facilitate Pi-scavenging by Pi-starved plants (Ghahremani et al., 2019). Although such observations need further investigation, it is conceivable that RXR1/RXR2, through interaction with other proteins, modulates other plant responses to P-limitation at the post-translational level (cf. Fig. 8). The interacting proteins At2g46520 and At1g64440, encoding exportin-2 and UDP-glucose epimerase 4 (UGE4) respectively, are interesting as well. In eukaryotic cells, nuclear export of proteins and RNAs by exportins are regulated by GTPases (Nielsen, 2020), providing, in addition to the effect of RXR1 on expression of RH-specific genes, another clue as to why RXR1/RXR2 are also found in the nucleus. Some alleles of UDP-galactose synthesizing UGE4 are known for altered RH phenotypes (Schiefelbein and Somerville, 1990; Seifert et al., 2002; Nguema-Ona et al., 2006), but not for altered RH elongation growth. UDP-galactose synthesis by UGE4 is important not only for cell wall synthesis but also for the organization and function of endomembranes (Wang et al., 2015), especially the trans-Golgi network and early endosome in Arabidopsis root epidermal cells, which are a sorting station of the secretory and vacuolar pathways. This puts UGE4 on the same stage as Rab GTPases, which are established key regulators of intracellular vesicle trafficking (Nielsen et al., 2008; Woollard and Moore, 2008; Stenmark, 2009; Nielsen, 2020).

Numerous genes affecting RH initiation, growth and shape have been identified over the past ∼20 years in Arabidopsis (Salazar-Henao et al., 2016; Shibata and Sugimoto, 2019) and other species (Marzec et al., 2015; Kim and Dolan, 2016; Kim et al., 2017). However, mutations in relatively few genes, such as*PCaP2* (Kato et al., 2019), *Lotus japonicas ROOTHAIRLESS LIKE4/5*(*LRL4/5*) (Breuninger et al., 2016), *ROOT HAIR- SPECIFIC1* (*RHS1*) or *RHS10* (Won et al., 2009), *GT-2-LIKE1* (*GTL1*) and its homolog *DF1* (Shibata et al., 2018) resulted in longer RHs. Similar to *rxr1* and *rxr2* RHs, *pcap2* RHs elongate at a higher rate, resulting in RHs that are ∼50% longer than wild-type RHs (Kato et al., 2019).*RHS1* and *RHS10* were identified *in silico* as genes harboring RH-specific *cis*-elements in their promoters (Won et al., 2009). *RHS10* encodes a proline-rich receptor-like kinase that localizes to the plasma membrane and exhibits association with RH cell walls (Hwang et al., 2016), which is very different to RXR1’s localization. Moreover, *rhs10* mutant RHs elongate at the same rate as wild-type RHs, but the tip-growing period is extended, resulting in ∼35% longer RHs, whereas *rxr1* and *rxr2* RHs elongate faster and ultimately are 75 to >100% longer than wild-type RHs (Fig. 3B, Fig. 5G). Furthermore, auxin or ethylene do not rescue RHS10–inhibited RH growth (Hwang et al., 2016), but they still function in *rxr1* mutant and RXR1 over-expresser, again indicating a mechanistic difference. The trihelix TFs GTL1 and DF1 repress RH growth by direct binding to the *RSL4* promoter (Shibata et al., 2018). Similar to *rhs10* or *RSL4* OX (Yi et al., 2010; Hwang et al., 2016), but in contrast to *rxr1*, *rxr2* or *pcap2*, *gtl1df1* double mutant RHs also elongate at the same rate as wild-type RHs, but their tip-growing period is prolonged, resulting in ∼50% longer RHs. In summary, these results suggest that RHS10, GTL1 and DF1 are involved in termination of the tip-growth process, while RXR1/RXR2 and PCaP2 negatively affect the rate of tip-growth. The phenotypic similarity of *rxr1*, *rxr2* and *pcap2* RH growth may indicate a closer connection between RXR1/RXR2 and phosphoinositides, although a general role of RXR1/RXR2 in RH lipid metabolism, vesicle transport and/or peroxisomal energy production is also possible (see above).

Mutants without RHs show reduced Pi uptake and compromised growth on soils when Pi availability is limited (Gahoonia et al., 2001; Brown et al., 2012; Tanaka et al., 2014). The same was observed with *RXR1* over-expressers in Arabidopsis and *Brachypodium*, which had substantially shorter RHs, lower Pi contents and reduced biomass irrespective of medium type or P-supply (Fig. 3, Table S2; Fig. 7). *Vice versa*, common bean genotypes with longer RHs have significantly higher Pi acquisition and shoot biomass (Miguel et al., 2015), similar to our results with the Arabidopsis *rxr1* mutant (Fig. 3, Table S2). Interestingly however, in transgenic *Brachypodium* lines constitutively overexpressing RSL2 or RSL3 bHLH TFs there was no consistent relationship between long RHs, increase in Pi uptake and higher biomass (Zhang et al., 2018). This led to the conclusion that increasing RH length through biotechnology can improve P uptake efficiency only if pleotropic effects, caused by transgene insertions or associated genomic rearrangements, on plant biomass are avoided. Non-transgenic knockout of *RXR1* and potential functional homolog(s) may help in this regard, and may prove to be an effective approach to develop crops with longer RHs that are more resource-use efficient and can improve soil health.

## METHODS

### Plant material, growth and treatment

*Arabidopsis thaliana* (ecotype Col-0) seeds were sterilized according to (Ying et al., 2012) and placed on square plates containing half-strength Murashige and Skoog (MS) medium ± phosphate (Caisson Labs), supplemented with 1% (w/v) sucrose, 0.5 mM MES-KOH (pH 5.7) and solidified with 0.4% (w/v) Gelzan^™^ CM (Millipore Sigma). The plated seeds were stratified for three days at 4 °C in the dark before placing the plates vertically in a growth chamber (22°C, 120 µmol^-2^ s^-1^ light intensity, 16 hrs light and 8 hrs dark cycle). Macronutrient deprivation treatments were done as described previously (Scheible et al., 2004; Bläsing et al., 2005; MORCUENDE et al., 2007; Bielecka et al., 2015). In brief, Arabidopsis seedlings were grown in liquid medium for seven days, transferred to nutrient-deficient liquid medium and harvested after another two days. For P-resupply, P-deprived seedlings were provided with 3 mM Pi and harvested 3 hrs later; cf. (MORCUENDE et al., 2007). Plant materials were harvested by rinsing twice in demineralized water, gently blotting on tissue paper and snap-freezing in liquid N2 before storage at -80°C until further use. The *rxr1* (SALK_048882), *rabd2c/rxr2* (SALK_054626), *lacs4* (CS65779 HM), *lacs6* (CS2103801), *at1g12000* (SALK_111594) and *at1g78850* (CS459880) T-DNA insertion mutant lines were obtained from ABRC. Homozygous mutant plants were identified using the protocol available at http://signal.salk.edu/.

### Identification and phylogenetic analysis of DUF506 proteins

To identify DUF506 proteins from different plant species, a BLASTPsearch was conducted on the Phytozome v12.1.6 website (https://phytozome.jgi.doe.gov/pz/portal.html) using the RXR1 domain 3 signature peptide (Figure 2) as query sequence. DUF506 protein sequenceswith low E value (<0.001) were downloaded and aligned using ClustalW online. Candidate proteins carrying the conserved motifs/domain (cf. Figure 2) were selected.For phylogenetic analysis, DUF506 protein sequences from *Arabidopsis thaliana*, *Brachypodium distachyon*, *Medicago truncatula*, and *Setaria viridis* were aligned and the phylogenetic tree generated using Geneious Prime software with default settings.

### Quantitative reverse transcription PCR (qRT-PCR)

Total RNA was isolated using the RNeasy^®^ plant mini kit (QIAGEN). cDNA was synthesized from 1 µg of DNase-treated RNA using SuperScript^™^ III reverse transcriptase (Invitrogen) according to manufacturer instructions. Quantitative PCR was performed using an Applied Biosystems 7900HT real-time PCR system with reaction conditions as follows: denaturation at 95°C for 10 min, followed by 40 cycles of 95°C for 15 s and 58°C for 1 min. Gene-specific primers were designed using PrimerQuest software available online (http://www.idtdna.com/Primerquest/Home/Index), and their sequences are listed in Table S1. Results were analyzed with SDS software v2.4 (Applied Biosystems). The Arabidopsis *GAPDH* (*At1g13440*) gene was used as reference (Czechowski et al., 2005). Experiments were repeated at least three times using cDNAs prepared from independent biological replicates.

### Measurements of root hair length

Seedlings were germinated vertically for 3 days on half strength MS plates containing 0.4% Gelzan™ CM (Millipore Sigma), and then transferred to plates containing either 0 (-P) or 675 µM (+P) phosphate, 10 or 100 nM IAA (Indole-3-acetic acid, Millipore Sigma), 100 nM or 1 µM ACC (1-aminocyclopropanecarboxylic acid, Millipore Sigma). Three days after transfer, root hair (RH) images were captured using a Nikon SMZ1500 fluorescence stereomicroscope with a scale bar of 1 mm. RH length was determined by an automatic measurement system in MATLAB. To that effect, root images were first transformed into white/black images, thereby canceling background noise and interference but keeping all RH details. Then the position and width of the primary root was determined, and the positions of RH tips detected. RH length was quantified by determining the distance between RH tip and the nearest position of the primary root. RH lengths were automatically calculated on a High Performance Computer (HPC). This RH measurement system was presented previously at Phenome 2018 (Tucson, AZ; https://community.plantae.org/article/5044038206316611401/phenome-2018-speaker-and-poster-abstracts). At least 700 RHs, located 2-6 mm from the tip of the primary root of 8-10 individual plants from each line or condition, were measured. The correctness of automatic measurements was checked by comparison to manual measurements of 50 randomly selected root hairs. Statistical analysis was done with the statistical toolbox in MATLAB using one way ANOVA and Tukey’s tests.

### Generation of over-expresser and complementation lines

Full-length coding regions of *At3g25240/RXR1* or *Bd2g58590* were amplified from *Arabidopsis* and *Brachypodium* seedling cDNA, respectively, using Phusion^®^ High- Fidelity DNA polymerase and cloned into GATEWAY^®^ entry vector pENTR™/SD/D- TOPO^®^ (Invitrogen). Sequence-confirmed vector was used for recombination into destination vector pMDC83 (Curtis and Grossniklaus, 2003) or pANIC6B (Mann et al., 2012). For *rxr1* complementation, a 1,461bp PCR fragment containing the *RXR1* promoter and coding region was amplified and ligated into destination vector pMU64 (Dr. Wei Liu, Noble Research Institute)which is based on pPZP200 (Hajdukiewicz et al., 1994). For *rabd2c/rxr2* complementation, a 233-bp promoter fragment of the *RabD2c* gene and its 609-bp coding region were fused into pMU64 via Gibson assembly strategy (Gibson et al., 2009). For histochemical GUS analysis, a 588-bp promoter fragment of *RXR1* was amplified from Arabidopsis genomic DNA and cloned into vector pBGWFS7 (Karimi et al., 2002). For calcium oscillation analysis, the *UBQ10::G-CAMP3-GFP* construct was transformed into *rxr1* mutant (Kwon et al., 2018). Constructs were introduced into *Agrobacterium tumefaciens* strain GV3101 (for Arabidopsis) or AGL1 (for Brachypodium) using a freeze-thaw procedure and were transformed into Arabidopsis (Col-0, and mutants) by floral dipping (Zhang et al., 2006), or Brachypodium (BD21-3) by embryogenic calli infection (Alves et al., 2009). Transformants were selected on 0.8% (w/v) agar plates containing half-strength MS medium (Caisson Labs) and 25 µg mL^-1^ hygromycin (HYG, Omega Scientific).

### Histochemical β-glucuronidase (GUS) staining and GFP imaging

GUS activity was analyzed according to (Jefferson et al., 1987). Briefly, tissues were incubated in a GUS staining solution containing 100 mM sodium phosphate (pH 7.0), 1 mM EDTA, 0.05% (v/v) Triton X-100, 1 mM potassium ferricyanide/ferrocyanide, and 0.5 mg mL^-1^ X-glucuronide (Goldbio) at 37°C for 1 to 3 hrs. After removal of the staining solution, tissues were cleared in 70% (v/v) ethanol. Images were acquired using Nikon SMZ1500 Stereomicroscope. GFP fluorescence in *pRXR1*::*RXR1-GFP* plants was detected and imaged with a Leica TCS SP8 confocal laser-scanning microscope (Ex: 488 nm; Em 507 nm). Propidium iodide (PI, 1 µg mL^-1^) was applied to visualize the cell shape.

### ATH1 gene-chip expression analysis

7-day-old Arabidopsis root tissues were conducted to microarray analysis. Two biological replicates were performed for each sample. Total RNA was isolated using RNeasy^®^ plant mini kit (QIAGEN) following the manufacturer’s instructions. RNA was quantified and evaluated for purity using a Nanodrop Spectrophotometer ND-100 (NanoDrop Technologies) and Bioanalyzer 2100 (Agilent). 100 ng of total RNA was used for the expression analysis of each sample using the Affymetrix GeneChip^®^ Arabidopsis Genome ATH1 Array (Affymetrix). Probe labeling, chip hybridization and scanning were performed according to the manufacturer’s instructions for 3’ IVT PLUS Kit (Affymetrix). Data normalization between chips was conducted using RMA (Robust Multichip Average) (Irizarry et al., 2003) implemented in Expression Console (Affymetrix). Gene selections based on Associative T-test (Dozmorov and Centola, 2003) were made using MATLAB. In this method, the background noise presented between replicates and technical noise during microarray experiments was measured by the residual presented among a group of genes whose residuals are homoscedastic. Genes whose residuals between the compared sample pairs that are significantly higher than the measured background noise level were considered to be differentially expressed. A selection threshold of 2 for transcript ratios and a Bonferroni-corrected P value threshold of 2.19202E^-06^ were used (Dozmorov and Centola, 2003). The Bonferroni-corrected P value threshold was derived from 0.05/N in these analyses, where N is the number of probes sets (22,810) on the chip.

### Co-Immunoprecipitation

Co-Immunoprecipitation (Co-IP) experiments were performed using the GFP-Trap^®_^M kit (ChromoTek). Proteins were extracted by grinding roots of 10-day-old Arabidopsis seedlings expressing *pCAMV35S*::GFP or *pRXR1*::*RXR1-GFP* in liquid N2, followed by homogenization of the powdered material in two volumes of ice-cold buffer containing 25 mM HEPES-KOH (pH 7.5), 1 mM EGTA, 1 mM EDTA, 20% (v/v) glycerol, 10 mM MgCl2, 25 mMNaF, 0.1% (v/v) Triton X-100, 0.1% (w/v) polyvinylpolypyrrolidone (PVPP), 1 mM phenylmethylsulfonyl fluoride (PMSF, Millipore Sigma), and 10 µg mL^-1^ cOmplete^™^ EDTA-free protease inhibitor cocktail (Millipore Sigma). Homogenates were centrifuged at 4°C and 16,000g for 10 min, and the supernatant re-centrifuged for 5 min and filtered through a layer of Miracloth (Calbiochem). Clarified extracts were incubated with pre-washed GFP-Trap^®^_M magnetic beads on an end-to-end tube rotator at 4°C for 1 hr, and the beads subsequently washed according to manufacturer instructions. Bound protein was eluted with 200 mM glycine (pH 2.5) and immediately neutralized with 1 M Tris (pH 10.4). Composition of the eluates was analyzed by LC-MS at the Charles W. Gehrke Proteomics Center (University of Missouri).

### Bimolecular fluorescence complementation (BiFC) assay

The BiFC analysis was performed as previously described (Waadt et al., 2014; Kudla and Bock, 2016). Briefly, *RXR1* was fused to the N-terminal EYFP in the pSITE-nEYFP vector, whereas *RabD2c* were fused with the C-terminal part of EYFP in pSITE-cEYFP. These vectors were co-transformed into *N. benthamiana* through agro-infiltration (Li, 2011). 48 hours after infiltration, fluorescence was detected and imaged using a Leica TCS SP8 confocal laser-scanning microscope (Ex: 514 nm; Em 527 nm).

### Pull-down assays and Western blotting

For heterologous expression of *RXR1*, its full-length (FL) cDNA was sub-cloned into pMAL^™^-c5X vector (NEB) carrying an N-terminal MBP (Maltose Binding Protein) tag. A C-terminally truncated (residues 1 to 165; 1-165) and two N-terminally truncated (residues 127 to 245; and residues 166 to 245) *RXR1*versions were also constructed and sub-cloned into the pMAL^™^-c5X expression vector. For recombinant protein production, constructs were separately introduced into the *Escherichia coli* (Express Competent cells C2523H, NEB). The purification was performed using amylose resin (NEB) according to the manufacture’s instruction. Protein concentration was quantified using Pierce^™^ rapid gold BCA protein assay kit (Thermo Fisher Scientific), and finally stored in -80°C. For the generation of His6-RabD2c, full-length cDNA of *RabD2c* were ligated into a pRSETA vector (Thermo Fisher Scientific). The proteins were heterologously expressed in *E. coli* (One Shot^®^BL21DE3pLysS cells, Thermo Fisher Scientific), purified using Ni- NTA Agarose (QIAGEN) according to the manufacturer’s instruction, and stored as described above.

For semi *in vivo* pull-down assays, the proteins were extracted from 10-day-old Arabidopsis *pRXR1::RXR1-GFP* seedling root tissue as described above. Protein extract (1 mg) was incubated with His6-RabD2c (1 μg) and Ni-NTA Agarose (QIAGEN) for 1 h at 4°C. The beads were washed three times with phosphate buffered saline (PBS, Millipore Sigma). Proteins bound to beads were dissociated by incubation in 100 μL of Novex^™^ Tris-Glycine SDS sample buffer (2x, Thermo Fisher Scientific) at 85°C for 2 min. Proteins were separated by Novex^™^ 10-20% Tris-Glycine gel (Thermo Fisher Scientific) and transferred to a Immobilon-P PVDF membrane (0.45 μM, Millipore Sigma). Western blotting was performed using 5000-fold diluted monoclonal mouse anti-GFP antibody conjugated to horseradish peroxidase (HRP) (MiltenyiBiotec), and was visualized using the ECL™ Prime Western Blotting System (Millipore Sigma).

For *in vitro* pull-down assays, His6-RabD2c (1 μg) was pretreated at 22°C for 15 min in 200 μL of PBS buffer containing 1 mM of the non-hydrolysable GTP analogue GTP-γ-S (Millipore Sigma) or GDP (Millipore Sigma), before incubation with Ni-NTA Agarose (QIAGEN) for 30 min at 4°C. Then the mixture was incubated with MBP-RXR1 or its truncated versions (1 μg) on a rotator for another 2 h at 4°C.The resin was washed five times with PBS, and bound proteins were denatured, separated and transferred as described above. Western blotting was performed using 5000-fold diluted monoclonal mouse anti-MBP antibody conjugated to HRP (MiltenyiBiotec).

### In vitro GTPase activity assay

The assay was performed as described in (Pan et al., 2006). Inorganic phosphate released by GTPase activity was quantified using the EnzChek Phosphate Assay Kit (Invitrogen) by measuring the increase in absorbance at λ=360 nm (A360) caused by production of 2- amino-6-mercapto-7-methylpurine. All assays were replicated at least three times and representative results are shown.

### Graphs and statistical analysis

Graphs were produced and statistical analysis conducted using SIGMAPLOT v12.5 (SystatSoftware, Inc., San Jose, CA, USA) and one-way ANOVA.

### Accession Numbers

Sequence data from this article can be found in the Phytozome 12 online genomic resource (https://phytozome.jgi.doe.gov/pz/portal.html) under the following accession numbers: *AtRXR1 (At3g25240), AtRabD2c/AtRXR2 (At4g17530), Arabidopsis DUF506* gene family members *(At1g12030, At1g62420, At1g77145, At1g77160, At2g20670, At2g38820, At2g39650, At3g07350, At3g22970, At3g54550, At4g14620 and At4g32480), AtRXR1* closest homologs from *Brachypodium distachyon (Bradi2g58590), Medicago truncatula (Medtr7g053310), Panicum virgatum (Pavir.8KG231400), Setaria viridis (Sevir.5G431800)* and *Triticum aestivum (TraesCS3B02G600900)*.

## SUPPLEMENTAL DATA

Supplemental Figure S1. Sashimi plots of selected, highly P-limitation-induced Arabidopsis gene transcripts.

Supplemental Figure S2. Expression of β-glucuronidase activity under control of the *AT3G25240* promoter.

Supplemental Figure S3. Conservation, phylogeny and distribution of DUF506 proteins in plants.

Supplemental Figure S4. Genotypes and their *RXR1/DUF506* expression levels.

Supplemental Figure S5. Higher frequency of [Ca^2+^] oscillations in *rxr1* mutant root hairs.

Supplemental Figure S6. Robust RH phenotypes of *RXR1* over-expresser or *rxr1* mutant in different growth conditions.

Supplemental Figure S7. Root hair length of *RXR1* over-expresser and *rxr1* mutant grown on Gelzan plates with or without phosphate addition.

Supplemental Figure S8. Relative abundance of root hair-specific gene transcripts in *rxr1* mutant and *RXR1* over-expresser.

Supplemental Figure S9. Effect of auxin and ethylene on RH length distribution, *RXR1* expression and hypocotyl length.

Supplemental Figure S10. Root hair length of *rabd2c/rxr2* mutant during P-limitation.

Supplemental Figure S11. Complementation of *rabd2c/rxr2* mutant.

Supplemental Figure S12. Overexpression of conserved RXR1 regions do not inhibit RH elongation growth.

Supplemental Figure S13. RH lengths of *Brachypodium* genotypes.

**Supplemental Table S1.** Sequences and purposes of primers used in the study.

**Supplemental Table S2.** P-contents and biomasses of wild type, *rxr1* mutant and *RXR1* over-expresser grown on different types of agar plates and P-supplies.

**Supplemental Table S3.** Relative expression of root-hair specific genes in root samples of *rxr1* mutant and *RXR1* over-expresser.

**Supplemental Table S4.** Expression of auxin and ethylene signaling-related genes in root samples of *rxr1* mutant and *RXR1* over-expresser.

**Supplemental Table S5.** Co-immunoprecipitated proteins and their mass-spectrometric peptide tags.

## ACKNOWLEDGMENTS

The authors wish to thank colleagues at Noble Research Institute for their help and support: Dr. Wei Liu, Alan Sparks, and Sunhee Oh for providing plasmid vectors, Dr. Pooja Pant for sharing RNA-Seq data, Sylvia Warner and Shulan Zhang for lab and greenhouse assistance, Amanda Hammon for plant care, Jianfei Yun and Dr. Qingzhen Jiang for providing Brachypodium transformation service, and Stacy Allen and Dr. Yuhong Tang for ATH1 transcript profiling service. We also thank Dr. Michael Udvardi for critical comments on the manuscript, and Dr. Brian Mooney (University of Missouri) for mass-spectrometric services. The work was funded by the Noble Research Institute LLC.

## AUTHOR CONTRIBUTIONS

S.Y. and W.-R.S. conceptualized and designed the research, analyzed and interpreted the data, and wrote the manuscript. S.Y. performed the experimental work. F.L. measured root hair lengths. E.B.B. measured root hair growth rates, recorded [Ca^2+^] oscillations and helped with or supervised some microscopic studies. All authors read and approved the final manuscript.

## Supplementary Information

**Figure S1.**
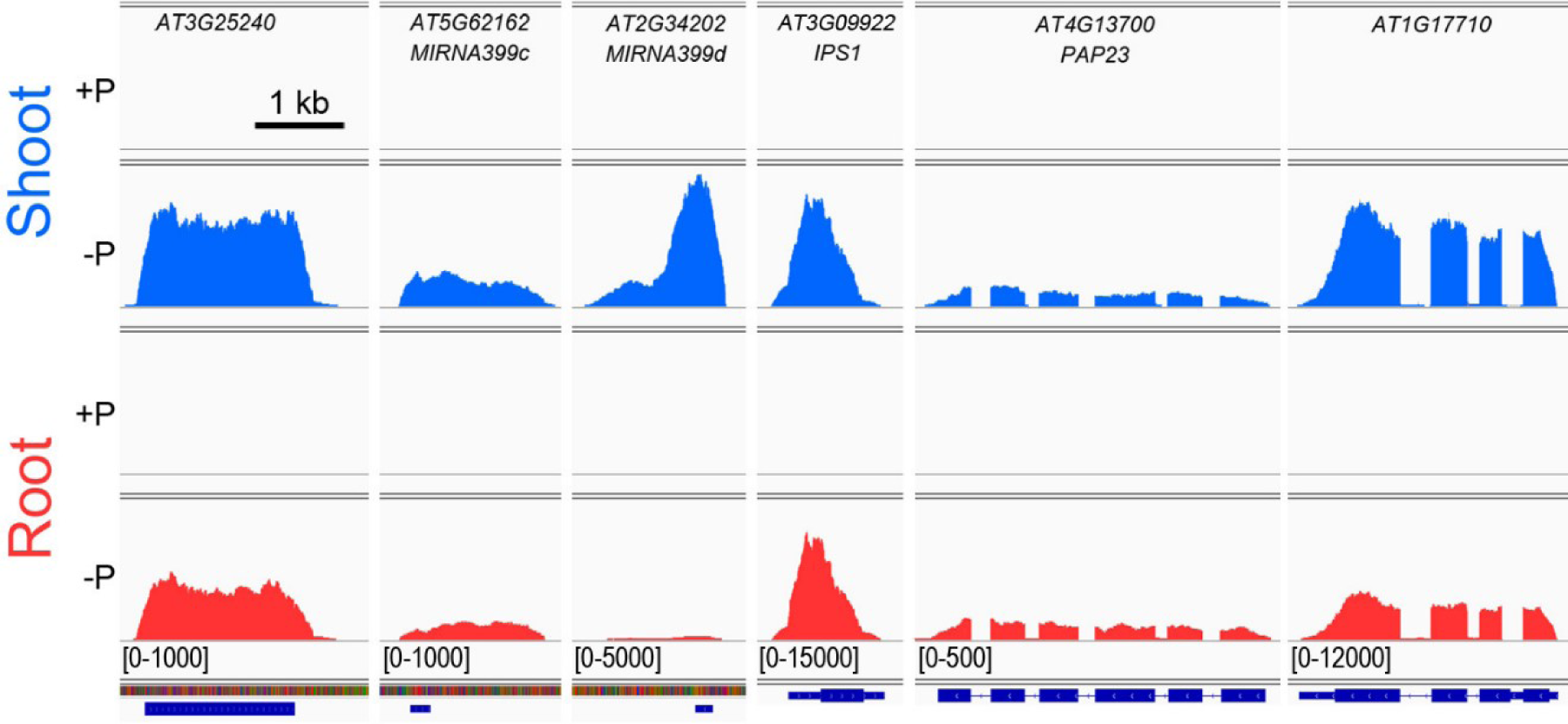
Sashimi plots of selected, highly P-limitation-induced Arabidopsis gene transcripts. RNA-seq read integrals are shown for (left to right): At3g25240 (RXR1), microRNA transcripts MIRNA399c and MIRNA399d, INDUCED BY PHOSPHATE STARVATION1 (IPS1), PURPLE ACID PHOSPHATASE23 (PAP23) and AT1G17710 (PEPC1) and were extracted from RNA-seq data generated from P-replete (+P) and P-limited shoots and roots of Arabidopsis plantlets. The height of an integral at a given position of a gene transcript indicates the number of times a nucleotide was sequenced. The range [0-XXXX] of the x-axis is the same for each of the four conditions, but differs between transcripts, as indicated. Gene transcripts are shown to scale, with a bar equal to 1 kb shown at the top left.

**Figure S2.**
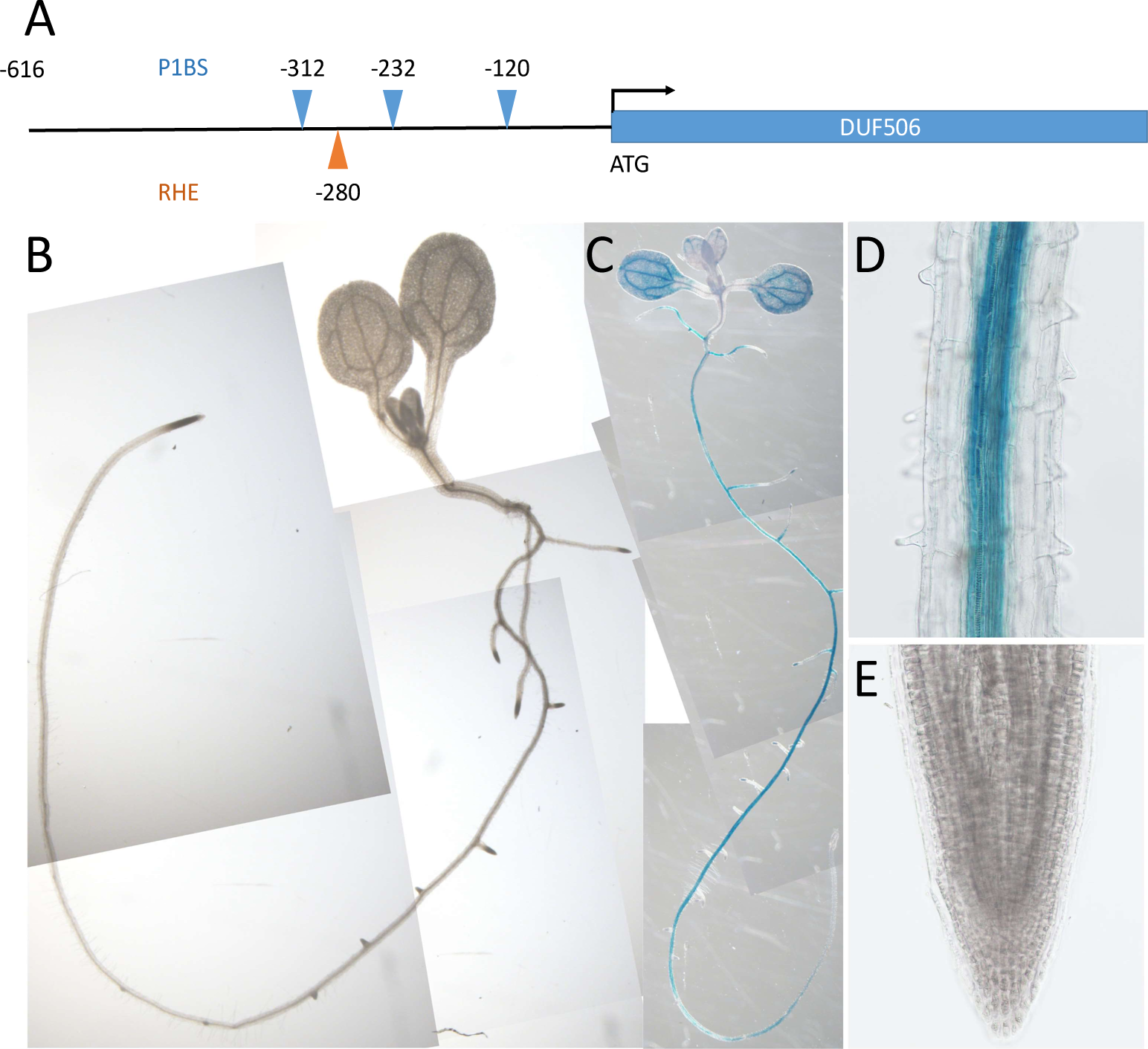
Expression of β-glucuronidase activity under control of the AT3G25240 promoter. (A)AT3G25240 promoter region with PHR1-binding sites (P1BS) and a root hair element (RHE), (B-E) Spatial β-glucuronidase activity in (B) P-replete and (C-E) P-deprived Arabidopsis seedlings. (D) and (E) show close-ups of the root-hair forming zone and the main root tip, respectively. Seedlings were germinated in +P liquid medium for 3 days before transfer to +P or –P liquid medium for another 3 days.

**Figure S3.**
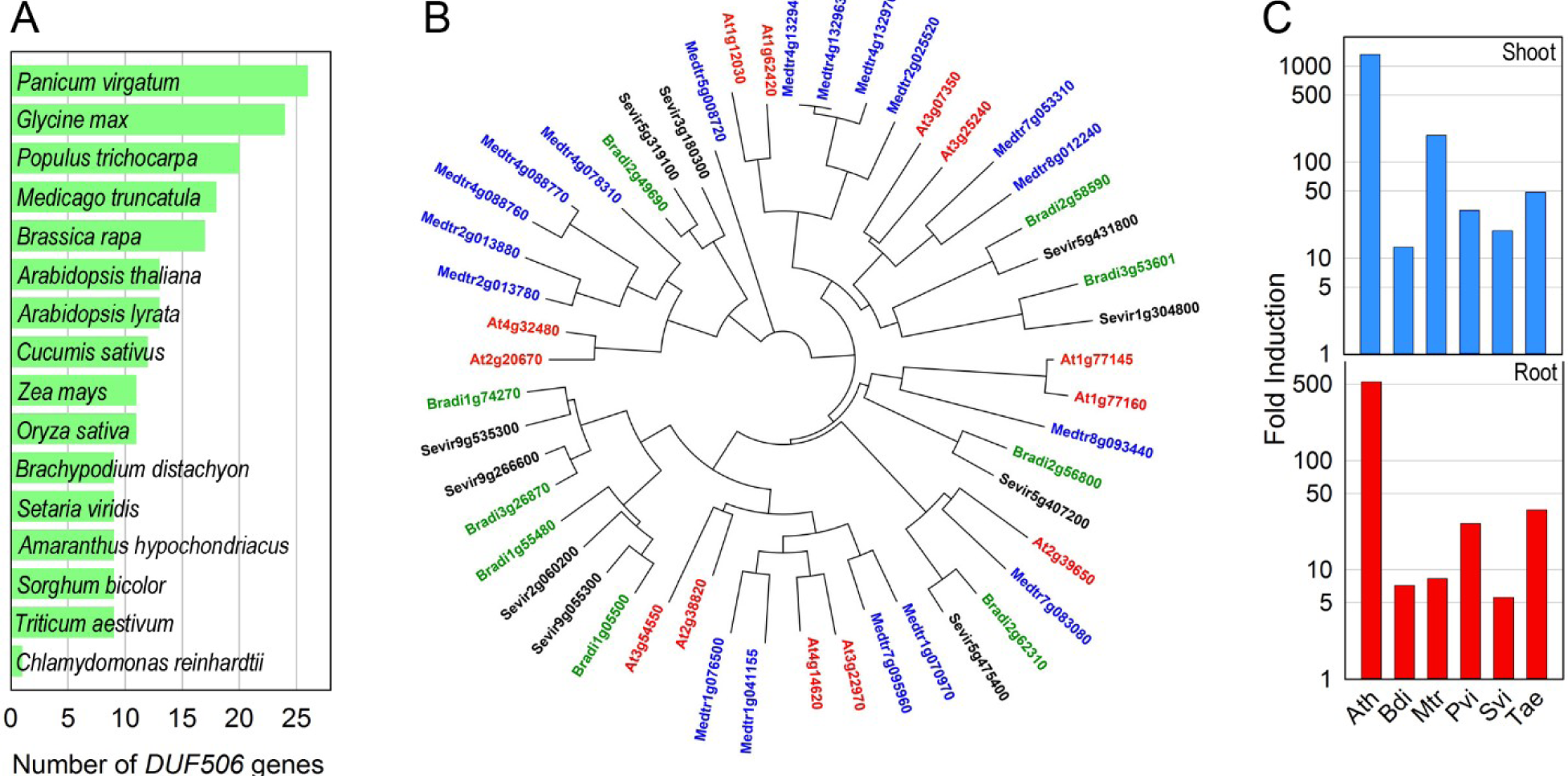
Conservation, phylogeny and distribution of DUF506 proteins in plants. (A) Number of DUF506 proteins found in fifteen plant species as well as the green algae Chlamydomonas reinhardtii. (B) Phylogeny of DUF506 proteins from Arabidopsis thaliana (red), Medicago truncatula (blue), Brachypodium distachyon (green) and Setaria viridis (black). (C) Induction of At3g25240 transcript and its closest homologs from Brachypodium distachyon (Bradi2g58590), Medicago truncatula (Medtr7g053310), Panicum virgatum (Pavir.8KG231400), Setaria viridis (Sevir.5G431800) and Triticum aestivum (TraesCS3B02G600900) during P-limitation in shoots and roots. The levels of induction are not expected to be quantitatively comparable as results from the different species originate from different experiments.

**Figure S4.**
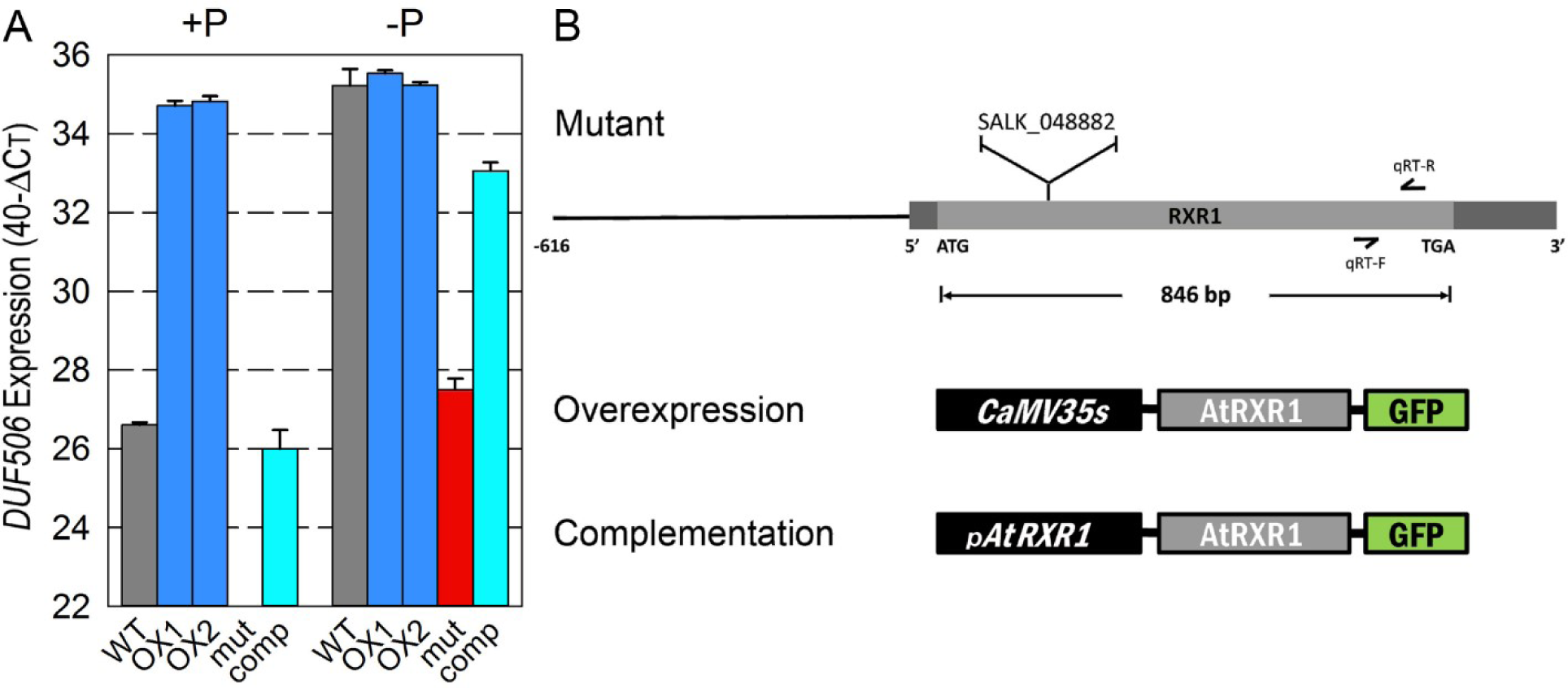
Genotypes and their RXR1/DUF506 expression levels. (A) RXR1/DUF506 expression levels in 6-day old seedlings of wild type, overexpression lines and T-DNA mutant (SALK_048882) in P-replete and P-limited conditions. (B) Schematic structure of T-DNA mutant (SALK_048882), overexpression and complementation constructs produced/used.

**Figure S5.**
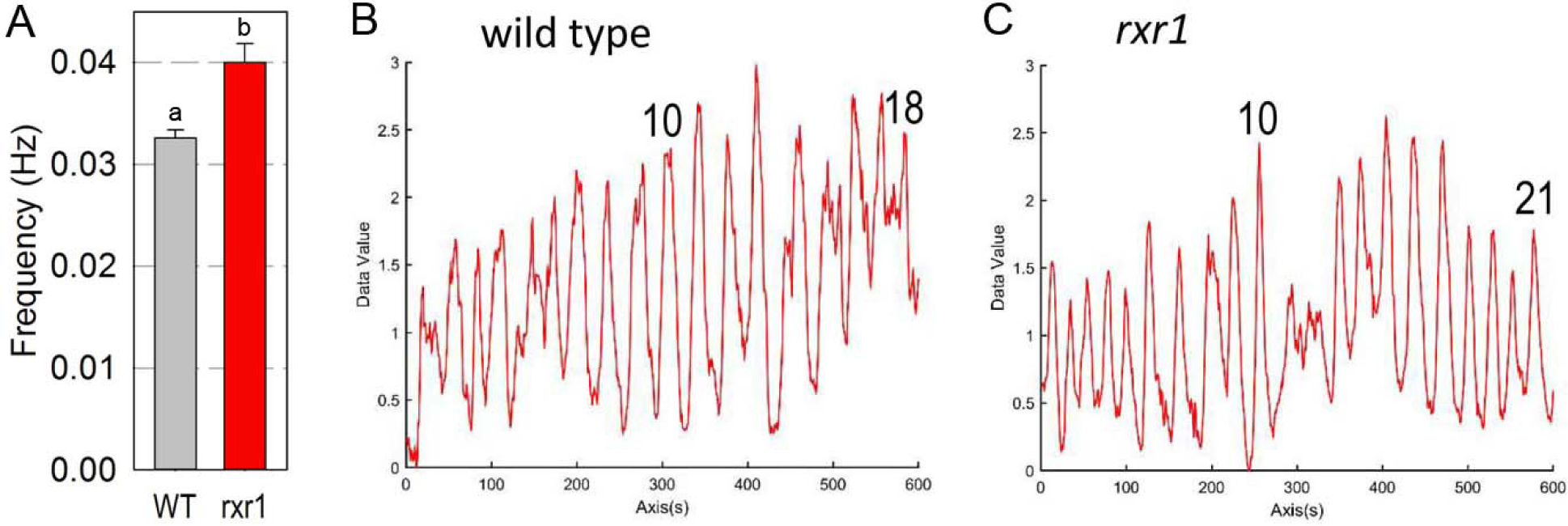
Higher frequency of [Ca^2+^] oscillations in rxr1 mutant root hairs. (A) Frequency of [Ca^2+^] oscillations in wild type (gray) and rxr1 mutant (red). The statistical significance is indicated (p < 0.01). (B, C) Example recordings of [Ca^2+^] oscillations in a root hair of (B) wild type and (C) rxr1 mutant.

**Figure S6.**
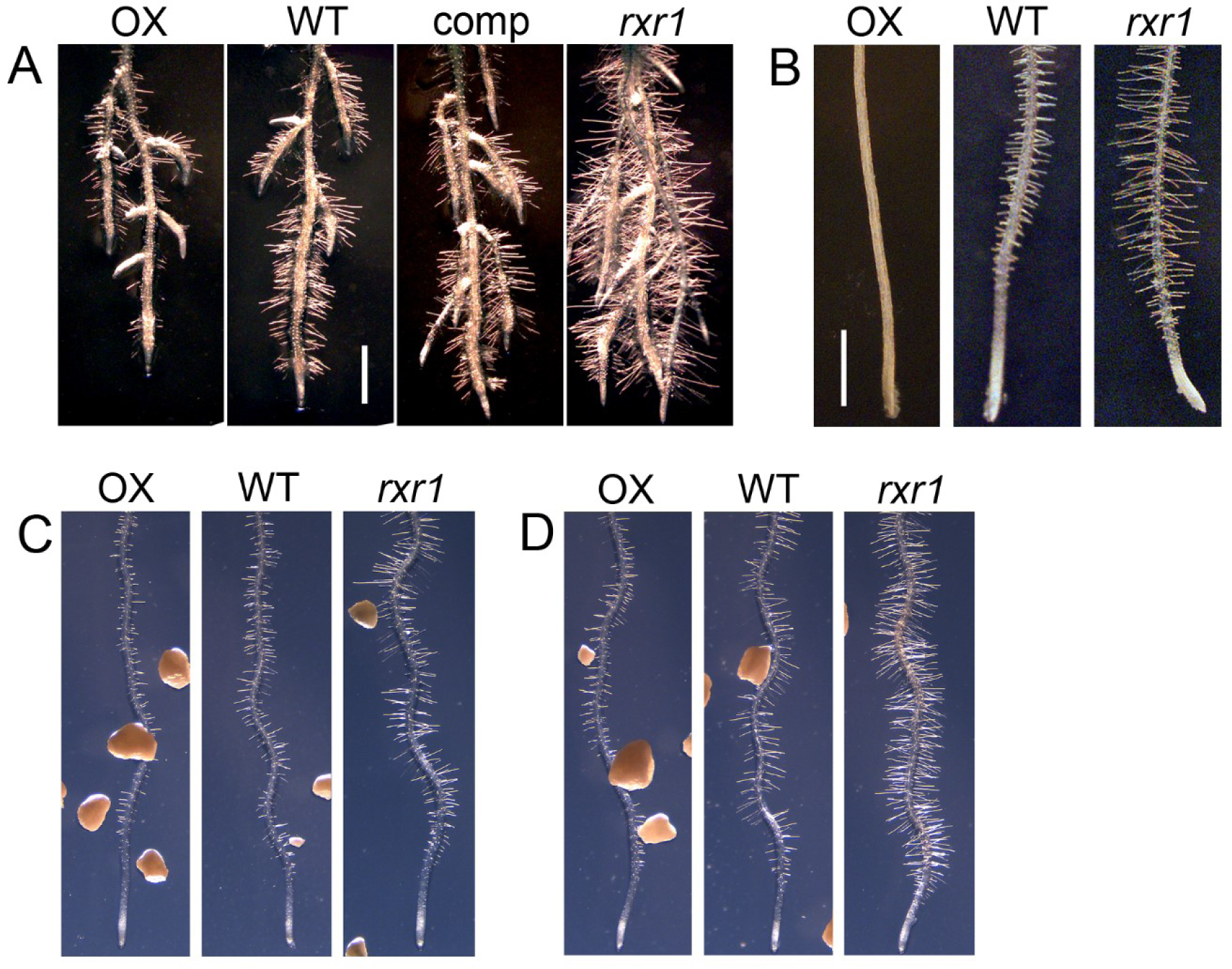
Robust RH phenotypes of RXR1 over-expresser or rxr1 mutant in different growth conditions. (A)Roots of 10-day old RXR1 over-expresser (OX), wild type (WT), rxr1 mutant (rxr1) and complemented rxr1 mutant (comp) directly sown and grown on agar (0.8% w/v) plates prepared with nutrient solution containing no phosphate. (B) Roots of 5-day old RXR1 OX, WT and rxr1 grown in liquid culture containing 675 µM phosphate. (C, D) Roots of 10-day old RXR1 OX, WT and rxr1 grown on vertical agar (0.8% w/v) plates containing phosphate-charged aluminum oxide beads providing steady concentrations of (C) 30 or (D) 8 µM phosphate (Hanlon et al., 2018; DOI: 10.1093/jxb/erx454). Bars = 1 mm.

**Figure S7.**
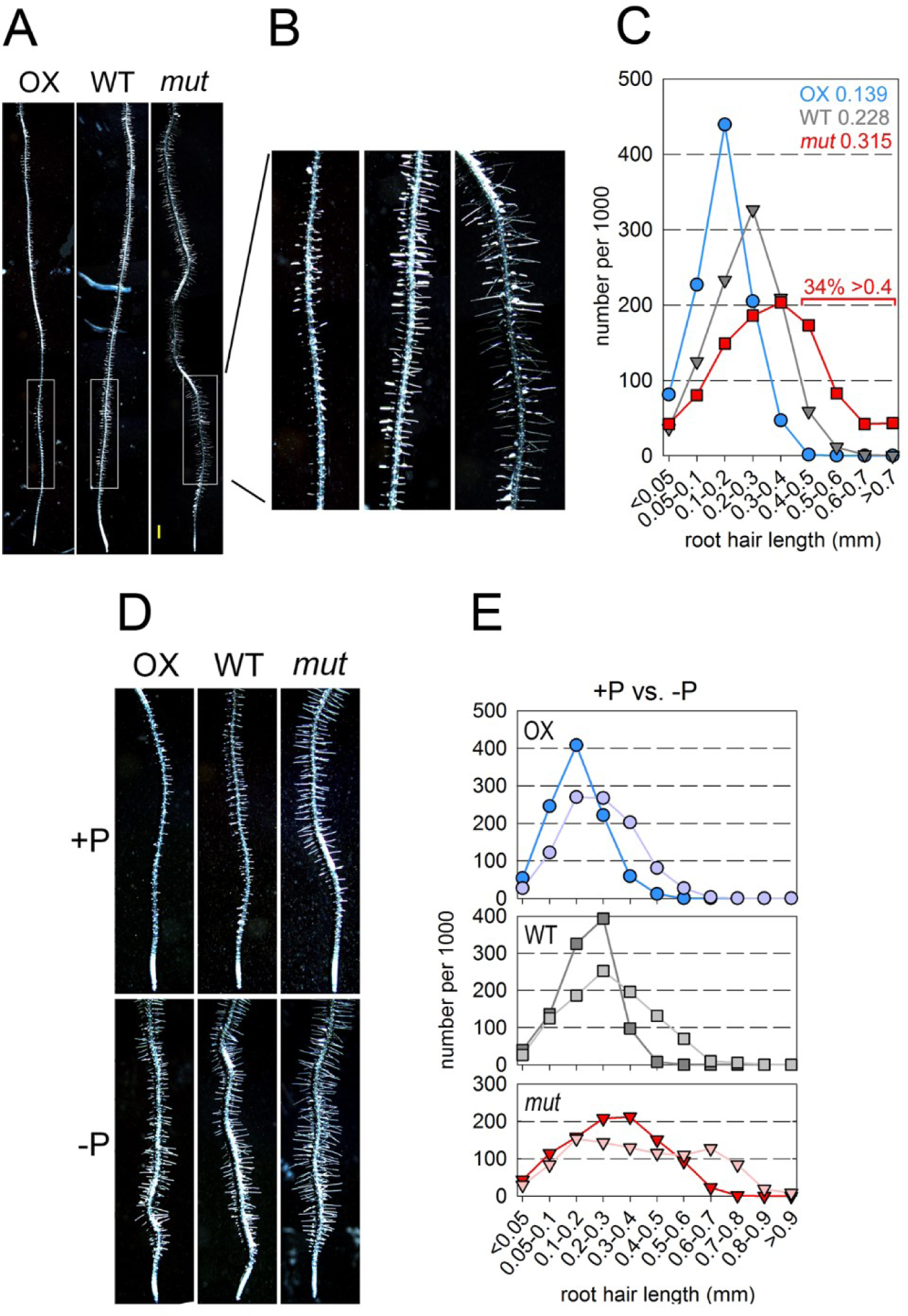
Root hair length of RXR1 over-expresser and rxr1 mutant grown on Gelzan plates with or without phosphate addition. (A) Roots of 6-day old RXR1 over-expresser (OX), wild type (WT) and rxr1 mutant grown for 3- days in liquid culture before transfer to Gelzan (0.4% w/v) plates prepared with nutrient solution containing 675 µM phosphate. (B) Magnification of the RH extension zones of the roots shown in (A). (C) RH length distribution of RXR1 over-expresser (OX), wild type (WT) and rxr1 mutant (mut) grown as described in the legend to (A). Between 787 and 815 root hairs were measured for each genotype and numbers for each length category normalized to 1000 RHs. Numbers in the top right corner of panel C indicate color-coded median values (in mm) for each genotype. (D) Roots of 6-day old plantlets grown as described in the legend to (A) with 675 µM phosphate (+P) or no phosphate (-P). (E) RH length distribution of RXR1 over-expresser (OX), wild type (WT) and rxr1 mutant (mut) grown as described in the legend to (D). Between 700 and 800 root hairs were measured for each genotype and numbers for each length category normalized to 1000 RHs.

**Figure S8.**
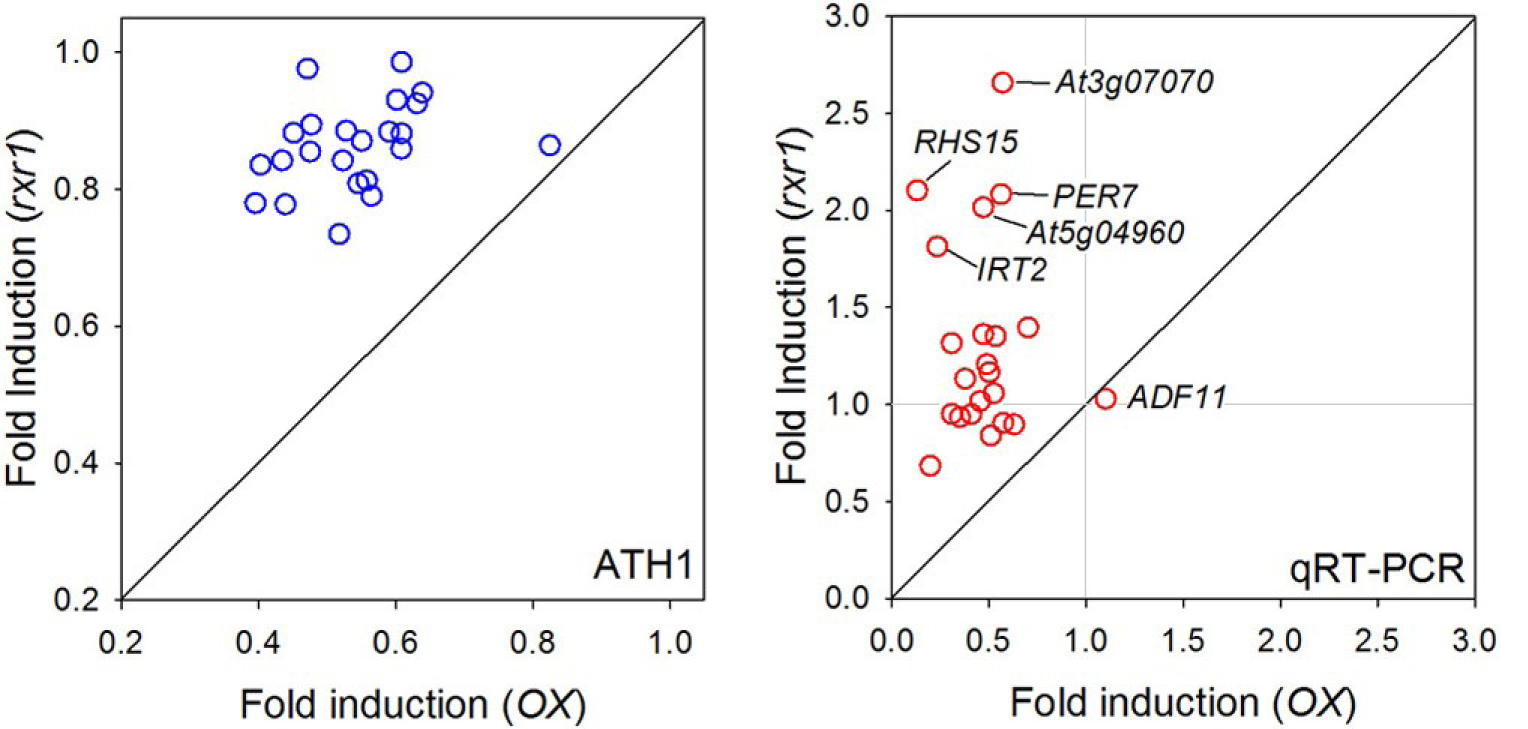
Relative abundance of root hair-specific gene transcripts in rxr1 mutant and RXR1 over-expresser. Fold induction of gene transcript abundances (rxr1 mutant vs. wild type, and RXR1 OX vs. wild type) are plotted against each other for ATH1 gene chip data (left panel, blue symbols) and qRT-PCR data (right panel, red symbols). Note that a “fold induction” of e.g. 0.5 equals a 2-fold decrease. Data for the depicted root hair-specific gene transcripts are also in Table S3, along with references.

**Figure S9.**
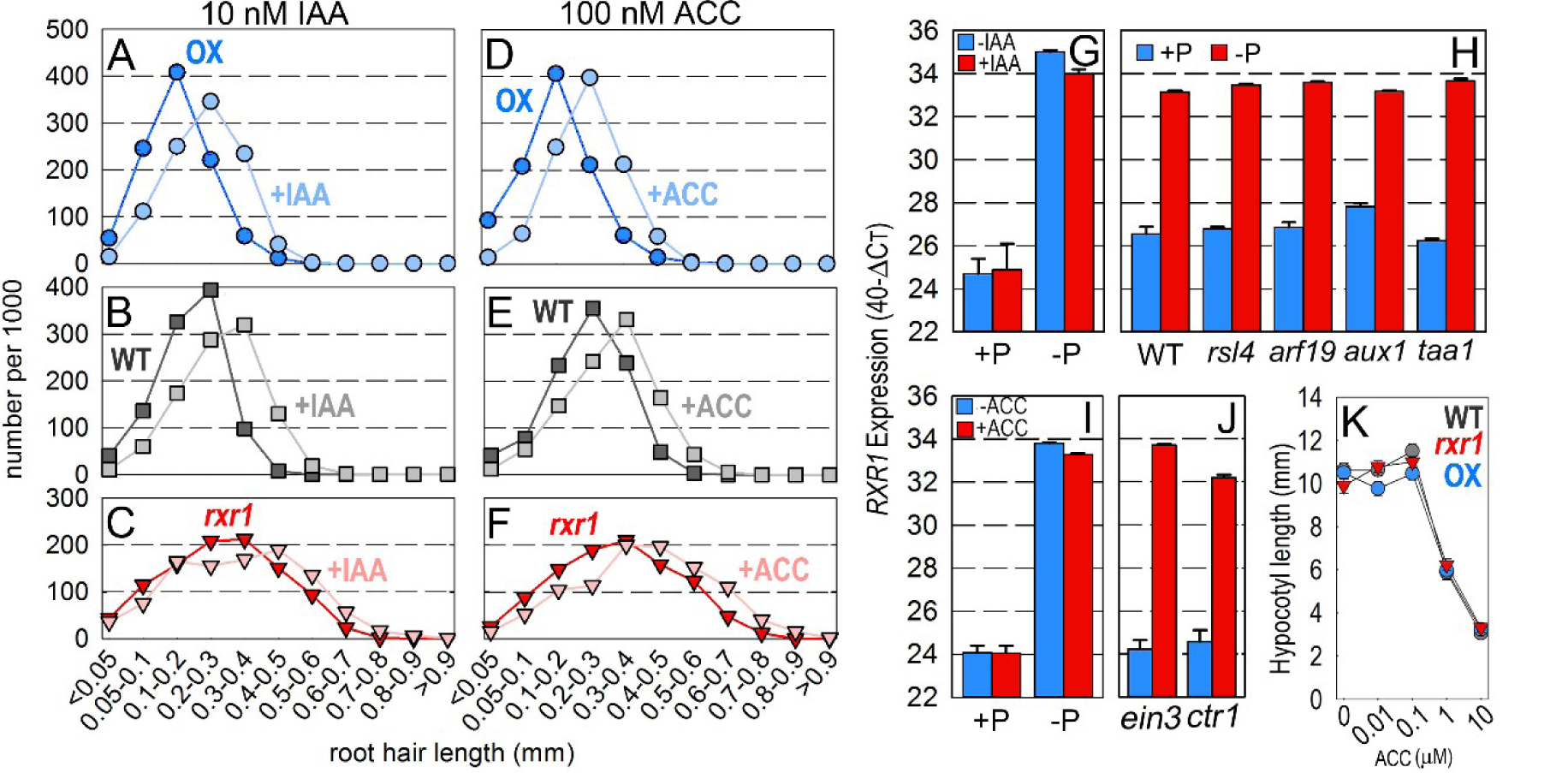
Effect of auxin and ethylene on RH length distribution, RXR1 expression and hypocotyl length. (A-F) RH length distribution plots for RXR1 over-expresser (OX), wild type (WT), and rxr1 mutant (rxr1) grown in the absence (darker colors) or presence of (A-C) 10 nM IAA or (D-F) 100 nM ACC (lighter colors). Seedlings were grown on Gelzan (0.4% w/v) plates prepared with nutrient solution containing 675 µM phosphate and phytohormone as indicated. For the distribution plots between 709 and 1241 root hairs were measured per genotype and treatment, and numbers for each length category were normalized to 1000 root hairs. (G-J) RXR1 expression, measured by qRT-PCR, (G) in the presence or absence of 100 nM IAA in P-replete or P-deprived conditions, (H) in wild type, rsl4, arf19, aux1 and taa1 mutants in P-replete or P-deprived conditions, (I) in the presence or absence of 1 µM ACC in P-replete or P-deprived conditions, and (J) in ein3 and ctr1 mutants in P-replete or P-deprived conditions.(K) Hypocotyl length (n=16) of RXR1 over-expresser (OX), wild type (WT), and rxr1 mutant (rxr1) in the presence of various ACC concentrations as described in Qu et al. 2007 (DOI: 10.1186/1471- 2229-7-3).

**Figure S10.**
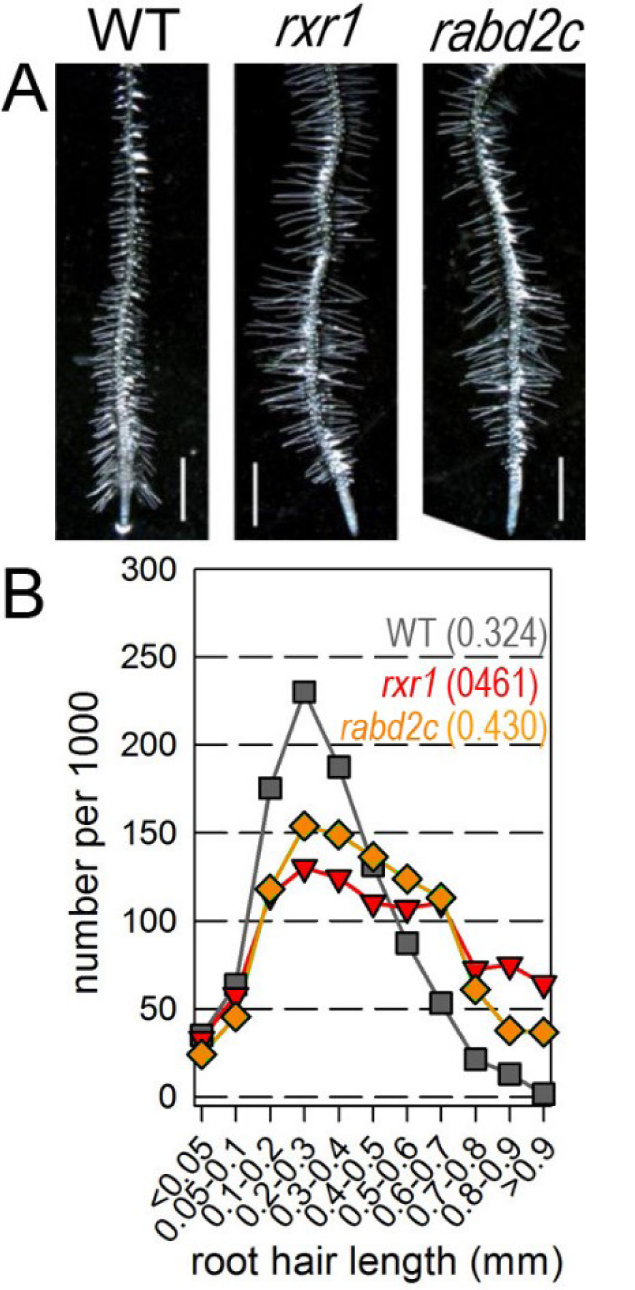
Root hair length of rabd2c/rxr2 mutant during P-limitation. (A) Roots of 6-day old wild type (WT), rxr1 mutant and rabd2c mutant grown for 3-days before transfer to Gelzan (0.4% w/v) plates prepared with nutrient solution containing no phosphate. Bar, 1 mm. (B) RH length distribution of wild type (gray squares), rxr1 mutant (red triangles) and rabd2c mutant (orange diamonds) grown as described in the legend to (A). Between 700 and 800 root hairs were measured for each genotype and numbers for each length category normalized to 1000 RHs. Numbers in parentheses indicate color-coded median values (in mm) for each genotype.

**Figure S11.**
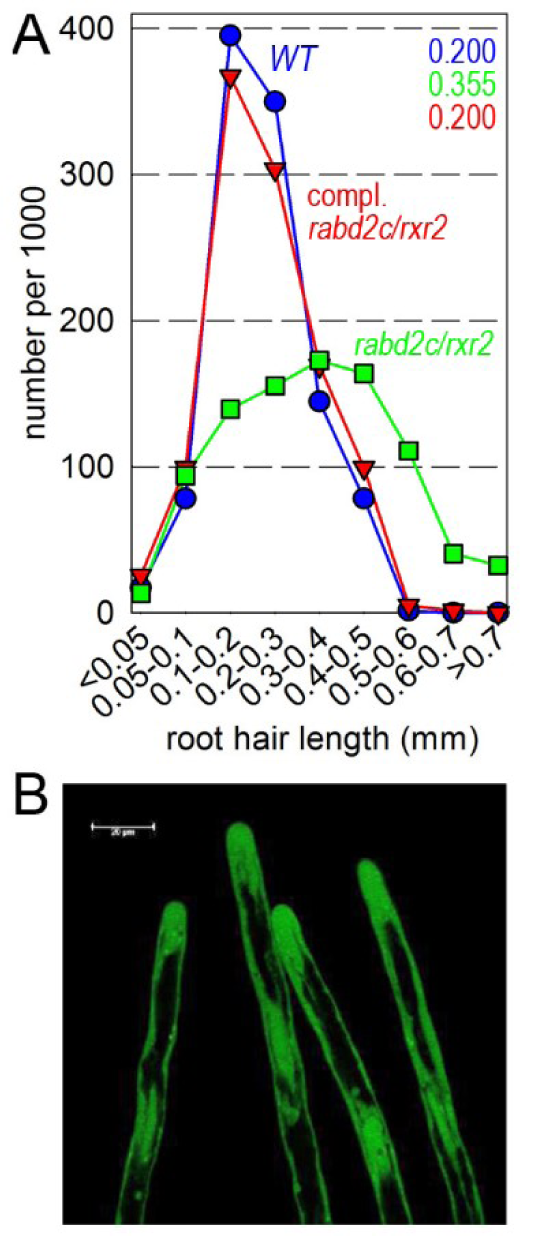
Complementation of rabd2c/rxr2 mutant. (A) Root hair length distribution of wild type (blue), rabd2c/rxr2 mutant (green) and complemented rabd2c/rxr2 mutant (red) grown on Gelzan (0.4% w/v) plates prepared with nutrient solution containing 675 µM phosphate. Between 776 and 804 root hairs were measured for each genotype and numbers for each length category normalized to 1000 RHs. Numbers in the top right corner indicate color-coded median values (in mm) for each genotype. (B) Expression in root hairs of RABD2c-GFP fusion protein expressed under control of the RABD2c promoter (bar = 20 µm).

**Figure S12.**
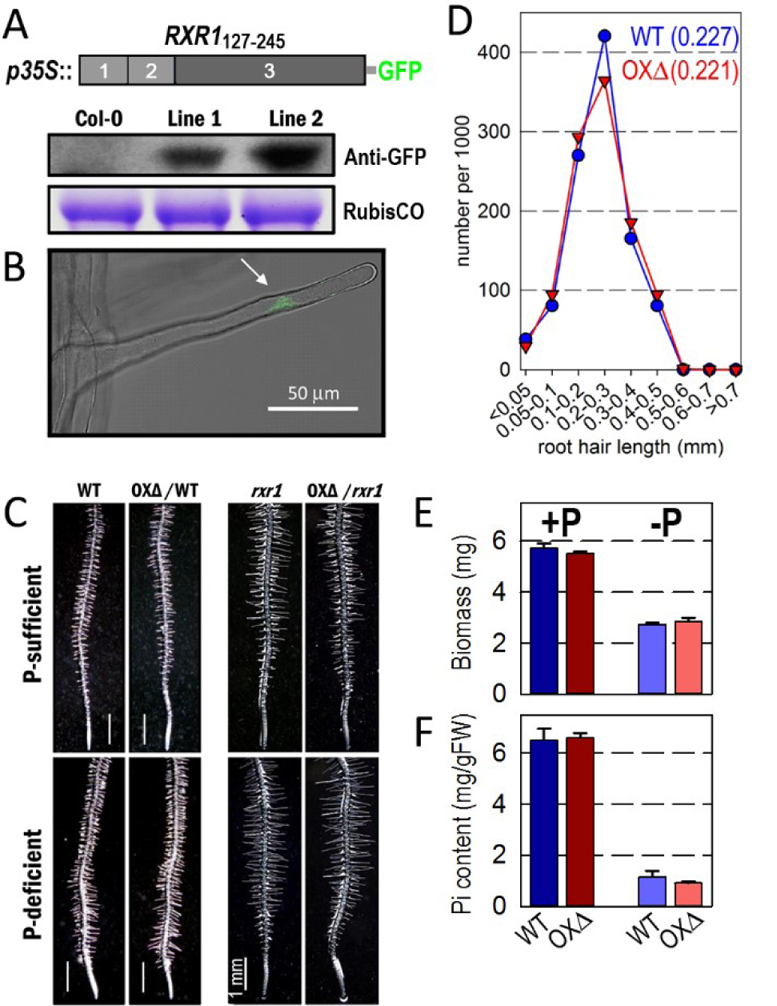
Overexpression of conserved RXR1 regions do not inhibit RH elongation growth. (A) Construct used for transformation of wild type and rxr1 mutant. The conserved motifs and domain that are sufficient for GTPase interaction and activity (cf. Figure 5) were fused to GFP driven by the CaMV-35S promoter (p35S). Western blotting confirms overexpression of RXR1_127-245_ -GFP fusion protein. (B) Expression of RXR1_127-245_ -GFP fusion protein in the RH nucleus. (C) RH phenotype of wild type and rxr1 mutant, both overexpressing RXR1_127-245_ - GFP, relative to the untransformed controls in P-sufficient and –deficient conditions. (D) RH length distribution of wild type (blue) and wild type overexpressing RXR1_127-245_ -GFP (OXΔ, red) grown on Gelzan (0.4% w/v) plates prepared with nutrient solution containing 675 µM or no phosphate for 3 days. Between 945 and 1047 RHs were measured for each genotype and numbers for each length category normalized to 1000 RHs. Numbers in the top right corner indicate color-coded median values (in mm) for each genotype. (E) Biomass and (F) phosphate (Pi) content of wild type (WT, blue) and wild type overexpressing RXR1_127-245_ -GFP (OXΔ, red) grown in P-sufficient (+P) and –deficient (-P) conditions. No statistical differences between wild type and OXΔ were revealed for biomass and Pi content. Plantlets used for (A), (E) and (F) and plantlets used for (B), (C) and (D) were 10 and 6-days old, respectively.

**Figure S13.**
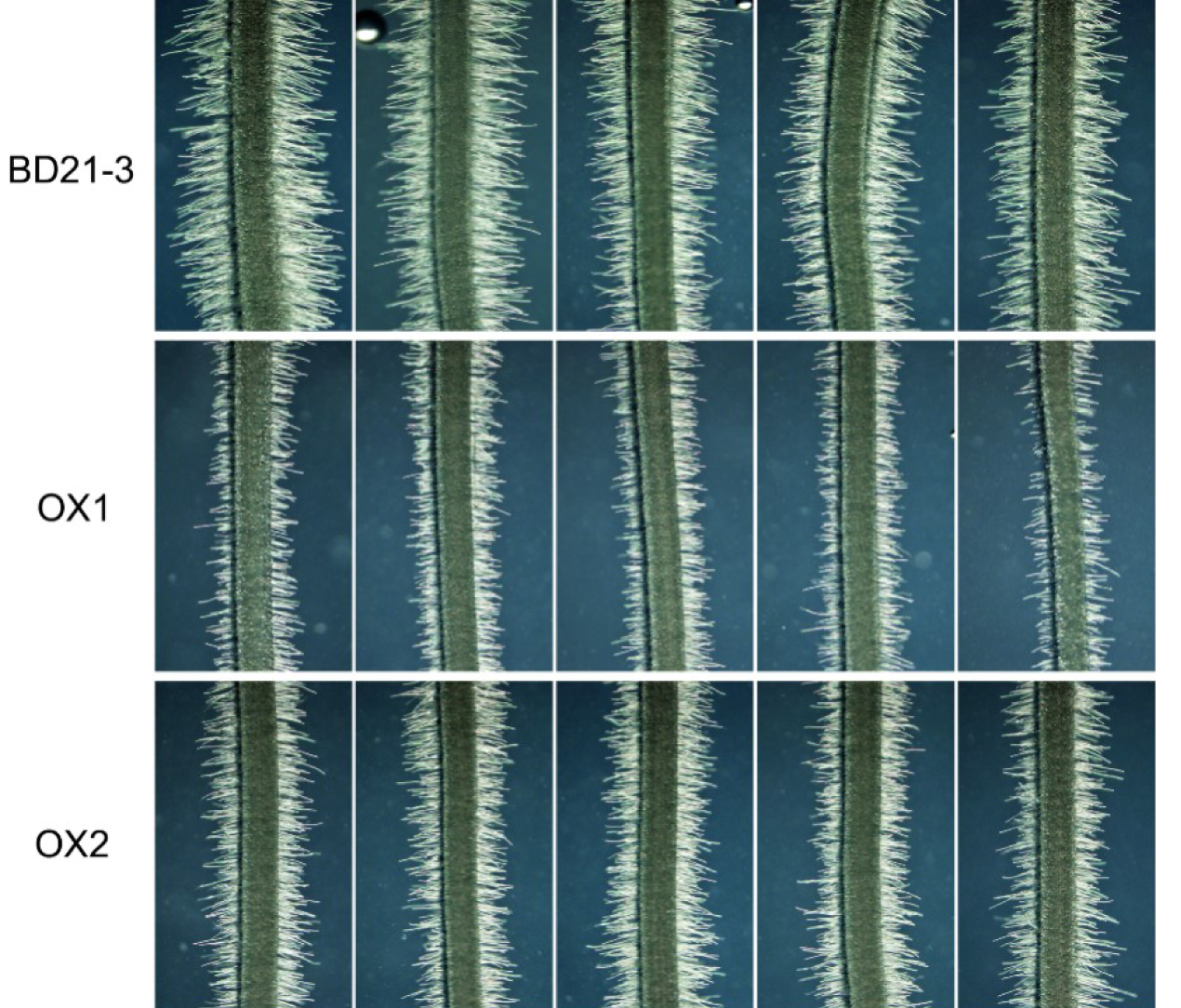
RH lengths of Brachypodium genotypes. Representative RH lengths of Brachypodium wild type (Bd21-3) (top) and Bd2g58590- overexpressing lines OX1 and OX2 (middle and bottom).

**Table S1.**
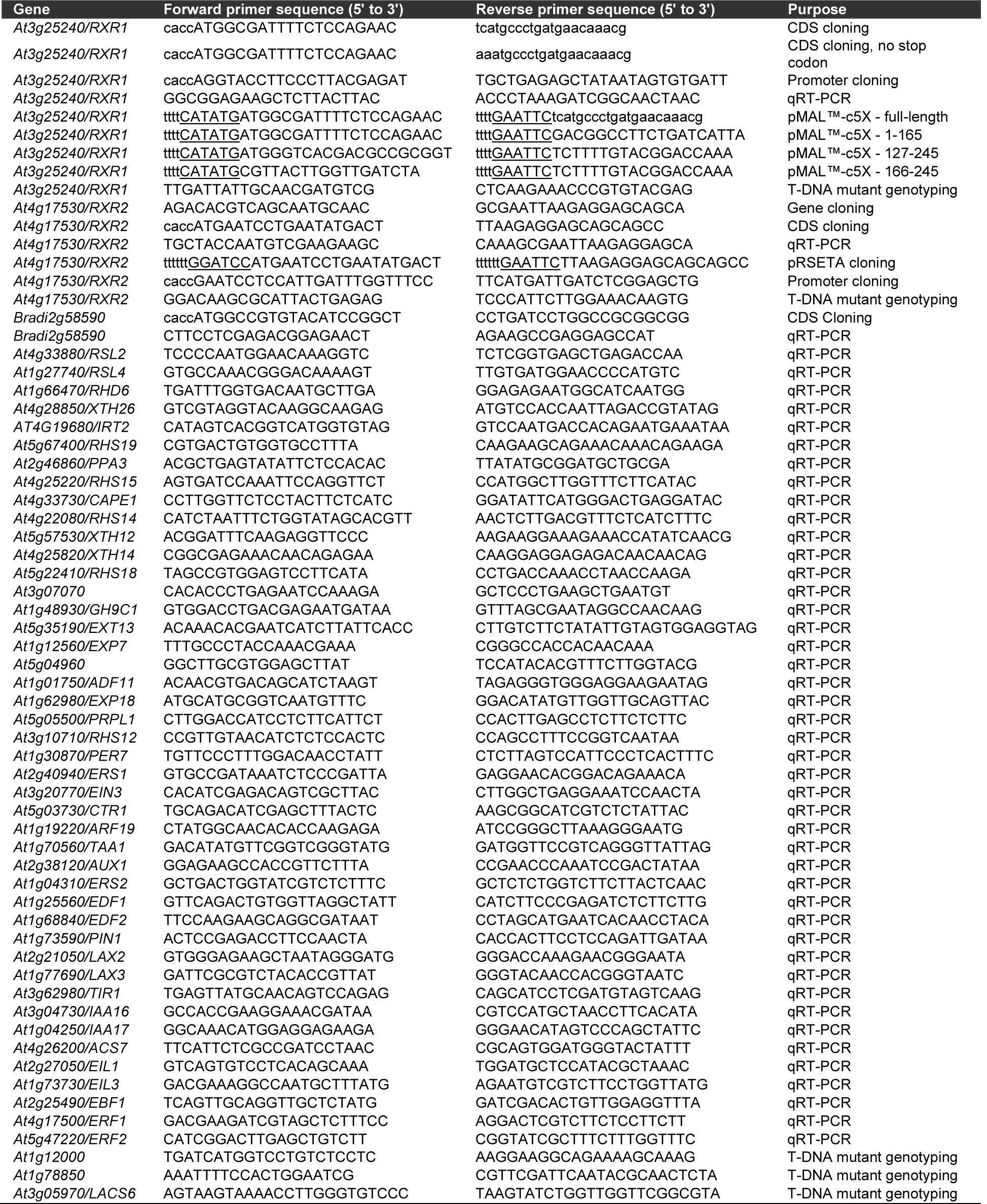
Sequences and purposes of primers used in the study.

**Table S2.**
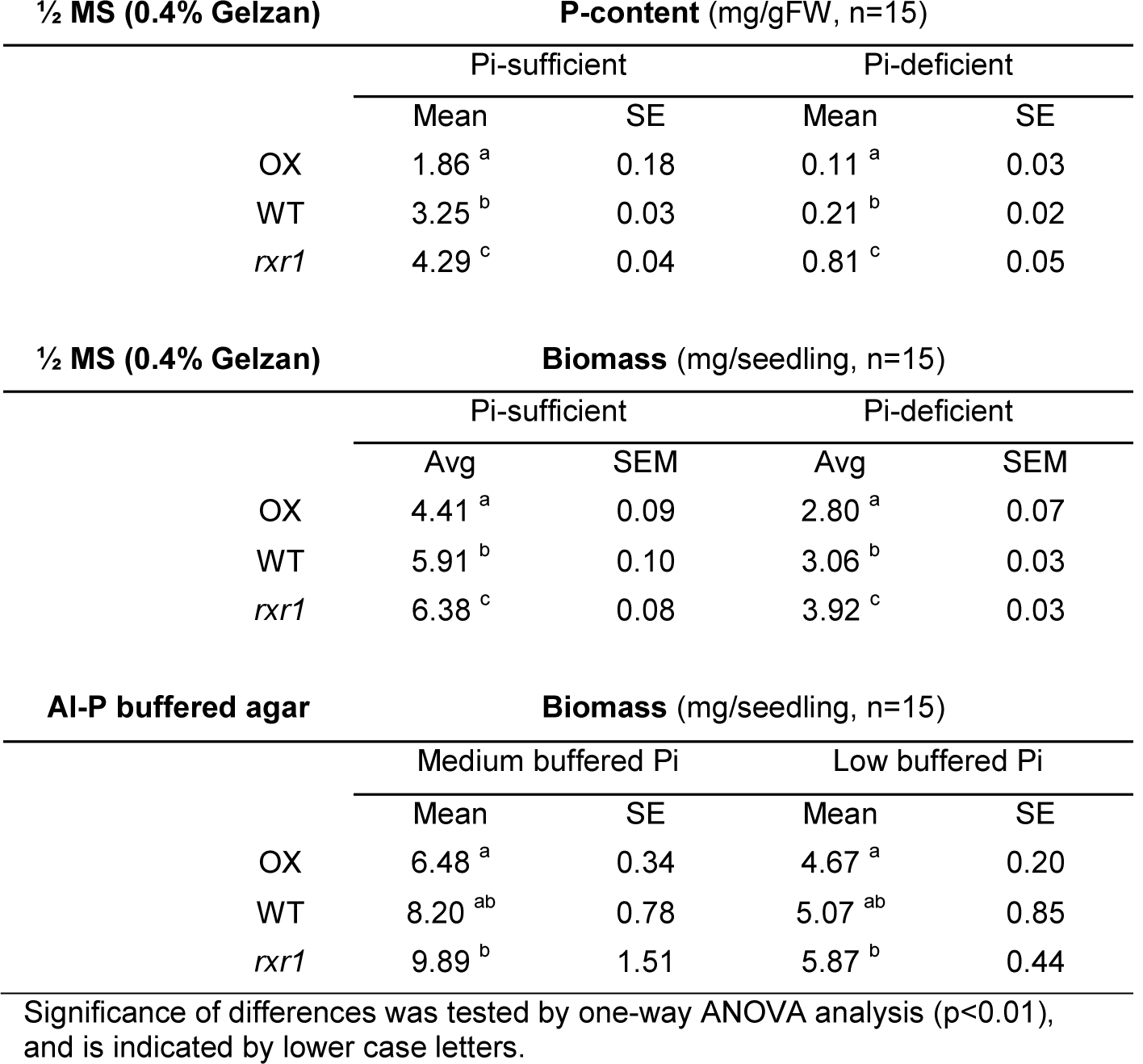
P-contents and biomasses of wild type, rxr1 mutant and RXR1 over-expresser grown on different types of agar plates and P-supplies.

| **½ MS (0.4% Gelzan)** |  | **P-content (mg/gFW, n=15)** |  |  |  |
| --- | --- | --- | --- | --- | --- |
|  |  | Pi-sufficient |  | Pi-deficient |  |
|  |  | Mean | SE | Mean | SE |
|  | OX | 1.86<sup>a</sup> | 0.18 | 0.11<sup>a</sup> | 0.03 |
|  | WT | 3.25<sup>b</sup> | 0.03 | 0.21<sup>b</sup> | 0.02 |
|  | *rxr1* | 4.29<sup>c</sup> | 0.04 | 0.81<sup>c</sup> | 0.05 |

| **½ MS (0.4% Gelzan)** |  | **Biomass (mg/seedling, n=15)** |  |  |  |
| --- | --- | --- | --- | --- | --- |
|  |  | Pi-sufficient |  | Pi-deficient |  |
|  |  | Avg | SEM | Avg | SEM |
|  | OX | 4.41<sup>a</sup> | 0.09 | 2.80<sup>a</sup> | 0.07 |
|  | WT | 5.91<sup>b</sup> | 0.10 | 3.06<sup>b</sup> | 0.03 |
|  | *rxr1* | 6.38<sup>c</sup> | 0.08 | 3.92<sup>c</sup> | 0.03 |

| **AI-P buffered agar** |  | **Biomass (mg/seedling, n=15)** |  |  |  |
| --- | --- | --- | --- | --- | --- |
|  |  | Medium buffered Pi |  | Low buffered Pi |  |
|  |  | Mean | SE | Mean | SE |
|  | OX | 6.48<sup>a</sup> | 0.34 | 4.67<sup>a</sup> | 0.20 |
|  | WT | 8.20<sup>ab</sup> | 0.78 | 5.07<sup>ab</sup> | 0.85 |
|  | *rxr1* | 9.89<sup>b</sup> | 1.51 | 5.87<sup>b</sup> | 0.44 |

Significance of differences was tested by one-way ANOVA analysis ( $p < 0.01$ ), and is indicated by lower case letters.

**Table S3.** Relative expression of root-hair specific genes in root samples of rxr1 mutant and RXR1 over-expresser.

| Gene ID | Ref. | Annotation | Fold change (relative to WT) |  |  |  | Fold change |
| --- | --- | --- | --- | --- | --- | --- | --- |
|  |  |  | ATH1 |  | qRT-PCR |  | ATH1 |
|  |  |  | OX | mutant | OX | mutant | RSL4 OX |
| At4g28850 | a-e | XTH26, xyloglucan endotransglucosylase/hydrolase 26 | 0.40 | 0.78 | 0.20 | 0.68 | 2.85 |
| At3g49960 | b-e | Peroxidase superfamily protein | 0.40 | 0.84 | 0.31 | 0.95 | nl |
| At4g19680 | a-e | IRT2, iron regulated transporter 2 | 0.43 | 0.84 | 0.23 | 1.81 | 2.42 |
| At5g67400 | abcef | RHS19, root hair specific 19 | 0.44 | 0.78 | 0.41 | 0.95 | 2.31 |
| At2g46860 | a-e | PPA3, pyrophosphorylase 3 | 0.45 | 0.88 | 0.46 | 1.02 | 1.61 |
| At4g25220 | abef | RHS15, root hair specific 15 | 0.47 | 0.98 | 0.13 | 2.10 | 2.29 |
| At4g33730 | abe | CAPE1, CAP-derived peptide 1 | 0.48 | 0.85 | 0.38 | 1.13 | nl |
| At4g22080 | abcef | RHS14, root hair specific 14 | 0.48 | 0.89 | 0.35 | 0.93 | 2.15 |
| At5g57530 | a-e | XTH 12, xyloglucan endotransglucosylase/hydrolase 12 | 0.52 | 0.73 | 0.51 | 0.84 | 2.29 |
| At4g25820 | a-e | XTH 14, xyloglucan endotransglucosylase/hydrolase 14 | 0.52 | 0.84 | 0.31 | 1.31 | 1.82 |
| At5g22410 | a-df | RHS 18, root hair specific 18 | 0.53 | 0.88 | 0.49 | 1.21 | 2.57 |
| At5g57540 | a-d | XTH 13, xyloglucan endotransglucosylase/hydrolase 13 | 0.55 | 0.81 | nd | nd | 1.77 |
| At3g07070 | abe | Protein kinase superfamily protein | 0.55 | 0.87 | 0.57 | 2.66 | 1.54 |
| At1g48930 | a-e | GH9C1, glycosyl hydrolase 9C1 | 0.56 | 0.81 | 0.54 | 1.35 | nl |
| At5g35190 | a-d | EXT13, extensin 13 | 0.56 | 0.79 | 0.70 | 1.40 | nl |
| At1g12560 | a-f | EXP7, expansin 7 | 0.59 | 0.88 | 0.57 | 0.90 | 2.7 |
| At5g04960 | a-d | Plant invertase/pectin methylesterase inhibitor superfamily | 0.60 | 0.93 | 0.47 | 2.01 | nl |
| At1g01750 | a-d | ADF11, actin depolymerizing factor 11 | 0.61 | 0.86 | 1.10 | 1.03 | nl |
| At1g62980 | a-f | EXP18, expansin 18 | 0.61 | 0.99 | 0.47 | 1.36 | 1.73 |
| At5g05500 | a-e | PRPL1/MOP10, proline-rich protein-like 1 | 0.61 | 0.88 | 0.53 | 1.06 | nl |
| At3g10710 | a-f | RHS12, root hair specific 12 | 0.63 | 0.92 | 0.50 | 1.16 | 2.18 |
| At1g30870 | b-e | PER7, peroxidase 7 | 0.64 | 0.94 | 0.56 | 2.08 | 1.57 |
| At1g27740 | b-d | RSL4, root hair defective 6-like 4 | 0.82 | 0.86 | 0.63 | 0.90 | 6.98 |

**Table S4.**
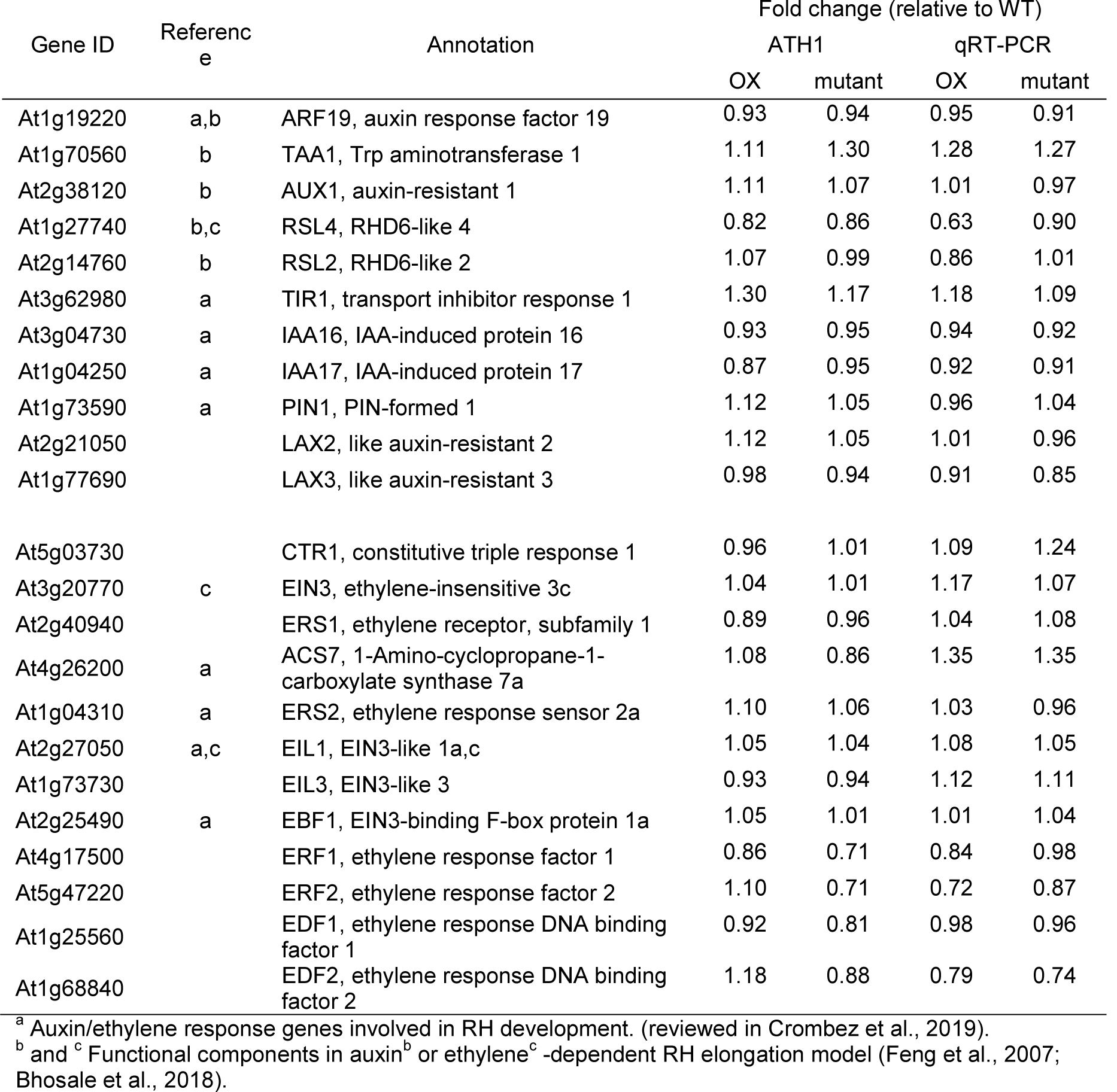
Expression of auxin and ethylene signaling-related genes in root samples of rxr1 mutant and RXR1 over-expresser.

| Gene ID | Reference | Annotation | Fold change (relative to WT) |  |  |  |
| --- | --- | --- | --- | --- | --- | --- |
|  |  |  | ATH1 |  | qRT-PCR |  |
|  |  |  | OX | mutant | OX | mutant |
| At1g19220 | a,b | ARF19, auxin response factor 19 | 0.93 | 0.94 | 0.95 | 0.91 |
| At1g70560 | b | TAA1, Trp aminotransferase 1 | 1.11 | 1.30 | 1.28 | 1.27 |
| At2g38120 | b | AUX1, auxin-resistant 1 | 1.11 | 1.07 | 1.01 | 0.97 |
| At1g27740 | b,c | RSL4, RHD6-like 4 | 0.82 | 0.86 | 0.63 | 0.90 |
| At2g14760 | b | RSL2, RHD6-like 2 | 1.07 | 0.99 | 0.86 | 1.01 |
| At3g62980 | a | TIR1, transport inhibitor response 1 | 1.30 | 1.17 | 1.18 | 1.09 |
| At3g04730 | a | IAA16, IAA-induced protein 16 | 0.93 | 0.95 | 0.94 | 0.92 |
| At1g04250 | a | IAA17, IAA-induced protein 17 | 0.87 | 0.95 | 0.92 | 0.91 |
| At1g73590 | a | PIN1, PIN-formed 1 | 1.12 | 1.05 | 0.96 | 1.04 |
| At2g21050 |  | LAX2, like auxin-resistant 2 | 1.12 | 1.05 | 1.01 | 0.96 |
| At1g77690 |  | LAX3, like auxin-resistant 3 | 0.98 | 0.94 | 0.91 | 0.85 |
| At5g03730 |  | CTR1, constitutive triple response 1 | 0.96 | 1.01 | 1.09 | 1.24 |
| At3g20770 | c | EIN3, ethylene-insensitive 3c | 1.04 | 1.01 | 1.17 | 1.07 |
| At2g40940 |  | ERS1, ethylene receptor, subfamily 1 | 0.89 | 0.96 | 1.04 | 1.08 |
| At4g26200 | a | ACS7, 1-Amino-cyclopropane-1-carboxylate synthase 7a | 1.08 | 0.86 | 1.35 | 1.35 |
| At1g04310 | a | ERS2, ethylene response sensor 2a | 1.10 | 1.06 | 1.03 | 0.96 |
| At2g27050 | a,c | EIL1, EIN3-like 1a,c | 1.05 | 1.04 | 1.08 | 1.05 |
| At1g73730 |  | EIL3, EIN3-like 3 | 0.93 | 0.94 | 1.12 | 1.11 |
| At2g25490 | a | EBF1, EIN3-binding F-box protein 1a | 1.05 | 1.01 | 1.01 | 1.04 |
| At4g17500 |  | ERF1, ethylene response factor 1 | 0.86 | 0.71 | 0.84 | 0.98 |
| At5g47220 |  | ERF2, ethylene response factor 2 | 1.10 | 0.71 | 0.72 | 0.87 |
| At1g25560 |  | EDF1, ethylene response DNA binding factor 1 | 0.92 | 0.81 | 0.98 | 0.96 |
| At1g68840 |  | EDF2, ethylene response DNA binding factor 2 | 1.18 | 0.88 | 0.79 | 0.74 |

<sup>a</sup> Auxin/ethylene response genes involved in RH development. (reviewed in Crombez et al., 2019).
<sup>b</sup> and <sup>c</sup> Functional components in auxin<sup>b</sup> or ethylene<sup>c</sup>-dependent RH elongation model (Feng et al., 2007; Bhosale et al., 2018).

**Table S5.**
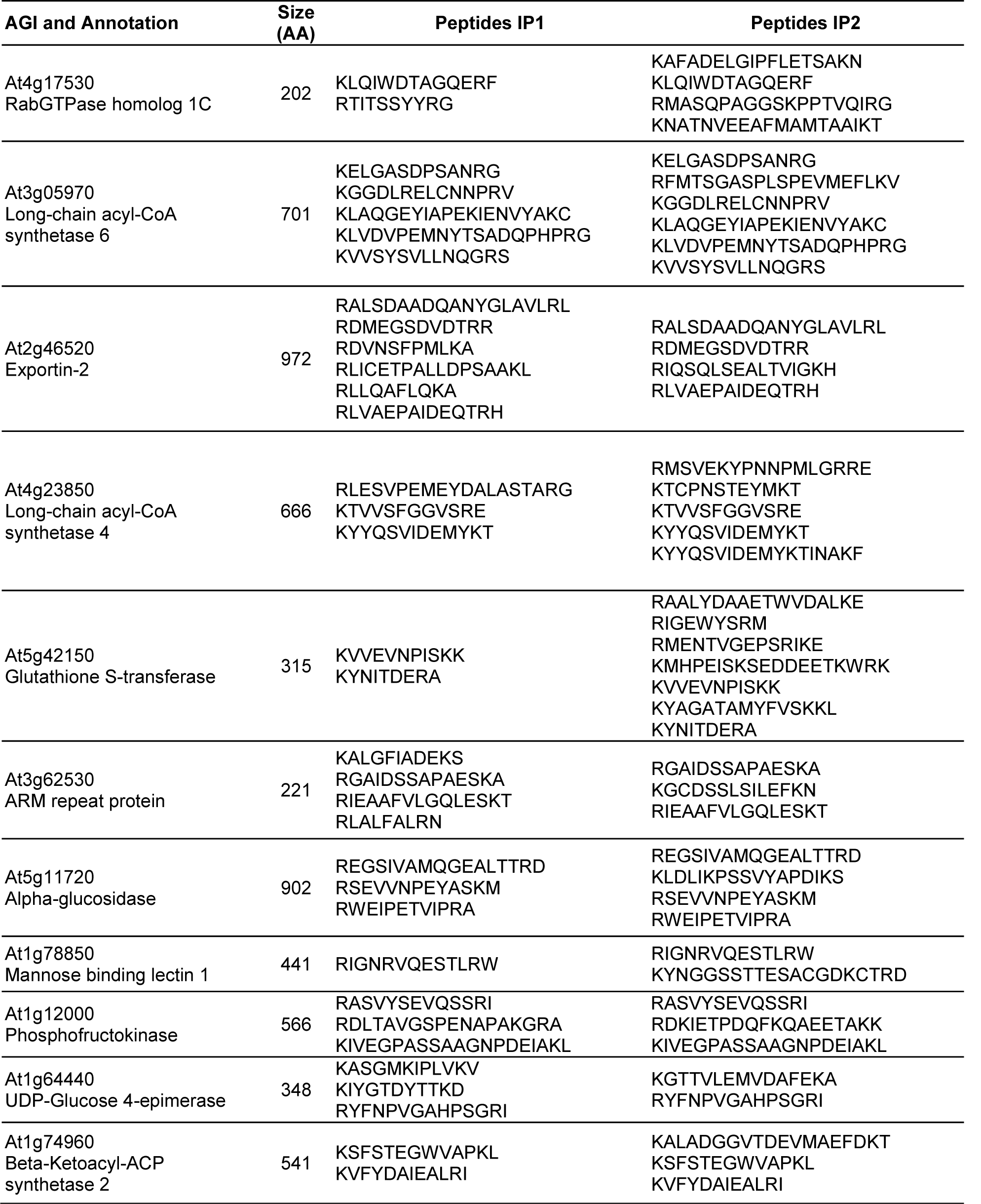
Co-immunoprecipitated proteins and their mass-spectrometric peptide tags.

